# An autoinhibitory clamp of actin assembly constrains and directs synaptic endocytosis

**DOI:** 10.1101/2020.03.06.981076

**Authors:** Steven J. Del Signore, Charlotte F. Kelley, Emily M. Messelaar, Tania Lemos, Michelle F. Marchan, Biljana Ermanoska, Markus Mund, Thomas G. Fai, Marko Kaksonen, Avital A Rodal

## Abstract

Synaptic membrane-remodeling events such as endocytosis require force-generating actin assembly. The endocytic machinery that regulates these actin and membrane dynamics localizes at high concentrations to large areas of the presynaptic membrane, but actin assembly and productive endocytosis are far more restricted in space and time. Here we describe a mechanism whereby autoinhibition clamps the presynaptic endocytic machinery to limit actin assembly to discrete functional events. We found that collective interactions between the *Drosophila* endocytic proteins Nwk/FCHSD2, Dap160/Intersectin, and WASp relieve Nwk autoinhibition and promote robust membrane-coupled actin assembly *in vitro.* Using automated particle tracking to quantify synaptic actin dynamics *in vivo*, we discovered that Nwk-Dap160 interactions constrain spurious assembly of WASp-dependent actin structures. These interactions also promote synaptic endocytosis, suggesting that autoinhibition both clamps and primes the synaptic endocytic machinery, thereby constraining actin assembly to drive productive membrane remodeling in response to physiological cues.

## Introduction

At neuronal presynaptic terminals, actin assembly affects many physiological processes including synapse morphogenesis, traffic of numerous vesicular cargoes, and synaptic vesicle endocytosis, organization, and mobility (Dillon and Goda, 2005; Nelson et al., 2013; Papandréou and Leterrier, 2018). However, the molecular mechanisms that control F-actin dynamics in space and time at presynaptic membranes are largely unknown. Presynaptic terminals maintain constitutively high local concentrations of actin-associated endocytic regulatory proteins at synaptic membranes (Reshetniak et al., 2020; Wilhelm et al., 2014), yet only a small fraction of this protein pool is likely to be active at any point in time (in response to vesicle release) and space (at <100 nm diameter endocytic sites), suggesting that the endocytic machinery is held in an inactive state at synaptic membranes. However, we do not know the mechanisms that maintain this machinery in an inactive state at the membrane, or how it is activated when and where it is needed.

One plausible mechanism to restrict membrane-cytoskeleton remodeling and endocytic activity to specific locations and times may lie in autoinhibition, which is a property of multiple endocytic proteins (Gerth et al., 2017; Kim et al., 2000; Rao et al., 2010; Stanishneva-Konovalova et al., 2016). One example is the F-BAR-SH3 protein Nervous Wreck (Nwk), which regulates synaptic membrane traffic at the *Drosophila* neuromuscular junction (NMJ) (Coyle et al., 2004; O’Connor-Giles et al., 2008; Rodal et al., 2008, 2011; Ukken et al., 2016) and whose mammalian homolog FCHSD2 regulates endocytosis and endocytic traffic in mammalian cells (Almeida-Souza et al., 2018; Xiao and Schmid, 2020; Xiao et al., 2018). Nwk/FCHSD2 proteins couple two activities: membrane remodeling and WASp-dependent actin polymerization (Almeida-Souza et al., 2018; Rodal et al., 2008; Stanishneva-Konovalova et al., 2016). Intramolecular autoinhibitory interactions between the Nwk F-BAR and its two SH3 domains mutually inhibit both Nwk membrane binding and activation of WASp (Stanishneva-Konovalova et al., 2016). Unlike other F-BAR-SH3 proteins, which are completely released from autoinhibition upon membrane binding (Guerrier et al., 2009; Meinecke et al., 2013; Rao et al., 2010), the SH3b domain of Nwk continues to restrict SH3a-mediated WASp activation even after Nwk binds membranes (Stanishneva-Konovalova et al., 2016). This suggests that autoinhibition allows Nwk-WASp to remain inactive even after recruitment to the membrane, thus keeping the endocytic machinery in a primed but inactive state. We hypothesized that additional binding partners of Nwk^SH3b^ may be required to fully activate membrane remodeling at discrete times and locations at the synapse.

An excellent candidate for release of Nwk autoinhibition at synapses is the endocytic adapter Intersectin (Dap160 in *Drosophila*). Intersectin interacts with numerous endocytic proteins to regulate endocytosis in mammalian cells (Henne et al., 2010; Okamoto et al., 1999; Praefcke et al., 2004; Pucharcos et al., 2000; Schmid et al., 2006; Sengar et al., 1999; Teckchandani et al., 2012), and has been implicated in several steps of the synaptic vesicle cycle (Evergren et al., 2007; Gerth et al., 2017; Jäpel et al., 2020; Pechstein et al., 2010, 2015). Of particular note, Intersectin recruits the Nwk homolog FCHSD2 to sites of endocytosis (Almeida-Souza et al., 2018), though it is not yet known how this affects FCHSD2 autoinhibition. In *Drosophila*, Dap160 interacts with WASp, Nwk, and other membrane remodeling proteins via its four SH3 domains (SH3A-D), and regulates the levels and localization of many of these proteins, including Nwk (Koh et al., 2004; Marie et al., 2004; Roos and Kelly, 1998). Further, *dap160* mutant phenotypes overlap with those of Nwk and WASp mutants, including impaired synaptic vesicle cycling and synaptic overgrowth (Coyle et al., 2004; Khuong et al., 2010; Koh et al., 2004; Marie et al., 2004). Finally, Intersectin and Dap160 shift localization from synaptic vesicle pools to the plasma membrane in response to synaptic activity (Evergren et al., 2007; Gerth et al., 2017; Winther et al., 2015), suggesting that Dap160 may provide the spatiotemporal link between salient physiological triggers and Nwk/WASp activation.

The high concentration and broad membrane distribution of inactive endocytic proteins (Reshetniak et al, 2020; Wilhelm et al, 2014) make it difficult to characterize the molecular dynamics of synaptic endocytosis. To overcome this barrier, we quantified discrete actin assembly events at the *Drosophila* NMJ as a proxy for productive endocytosis, as actin assembly is both a primary target of the endocytic apparatus under investigation, and is required for synaptic vesicle endocytosis in all forms, including at the *Drosophila* NMJ (Kononenko et al., 2014; Wang et al., 2010; Wu et al., 2016). This synapse is an ideal system to investigate the molecular dynamics of the endocytic machinery due to its large size, ease of genetic manipulation, and accessibility to live and super-resolution imaging. Here we combine *in vitro* biochemical approaches with quantitative imaging at the NMJ to define the interactions among Dap160, Nwk, and WASp that relieve autoinhibition. These interactions drive robust membrane-associated actin assembly *in vitro*, regulate the frequency and dynamics of synaptic actin structures *in vivo*, and are functionally required for normal endocytosis at the NMJ.

## Results

### Actin assembles in discrete dynamic patches despite broad distribution of presynaptic membrane-cytoskeleton remodeling machinery

While the importance of actin in synaptic endocytosis is clear (Kononenko et al., 2014; Wang et al., 2010; Wu et al., 2016), until now there has been no quantitative analysis of individual actin-dependent membrane-remodeling events at synapses. To better understand presynaptic F-actin dynamics and to identify sites where the cytoskeleton and membrane remodeling machinery is active, we quantified individual F-actin assembly events by spinning disk confocal microscopy of NMJs presynaptically expressing fluorescent actin probes. We performed these experiments under resting conditions, where vesicle release is spontaneous at a rate of ~5-6 vesicles/10 μm^2^/min (Akbergenova et al., 2018; Melom et al., 2013), presumably requiring a similar rate of compensatory endocytosis (Sabeva et al., 2017). We compared GFP::actin, a <u>G</u>FP-tagged <u>m</u>oesin F-<u>a</u>ctin binding domain (GMA), and Lifeact::Ruby. The predominant structures labeled by all of these markers at the presynaptic membrane were transient patches (**Movie 1, Fig 1A, Fig 1 S1A**), as has been previously observed (Nunes et al., 2006; Pawson et al., 2008; Piccioli and Littleton, 2014). We then quantified individual actin patch dynamics using automated particle tracking and quantification (Berro and Pollard, 2014), which captured on the order of 30-50% of visible actin structures (see methods for more detail). Imaging at 0.25Hz, we found an average of 1.2 GMA patches/10μm^2^/min, exhibiting a mean duration of 48.0 sec +/− 45.6 sec, with an average relative amplitude of 68% +/− 32% ((Imax-Imin)/Imean) (**Fig 1B-D**). Quantification of GFP::actin and lifeact::Ruby showed very similar dynamics to GMA, suggesting that these measurements robustly reflect the underlying actin dynamics and not the specific properties of a particular probe. As we noted a strong floor effect with short-duration patches, we also performed imaging at 1Hz, which could not capture the entire lifetime distribution due to photobleaching but was able to identify a larger population of short duration patches (**Fig 1S1B**) with an average duration of ~16 +/− 20 seconds. Given this range of measurements at different sampling frequencies and the efficiency of our automated detection, we estimate that patch frequency is between 2.8-10.3 events/10 μm^2^/min (see Methods for calculations), on par with the expected frequency of endocytic events, and with a similar albeit broader distribution of durations compared to yeast (15 seconds; Berro and Pollard, 2014) and mammalian cells (~40 seconds; Taylor et al., 2011).

**Figure 1.**
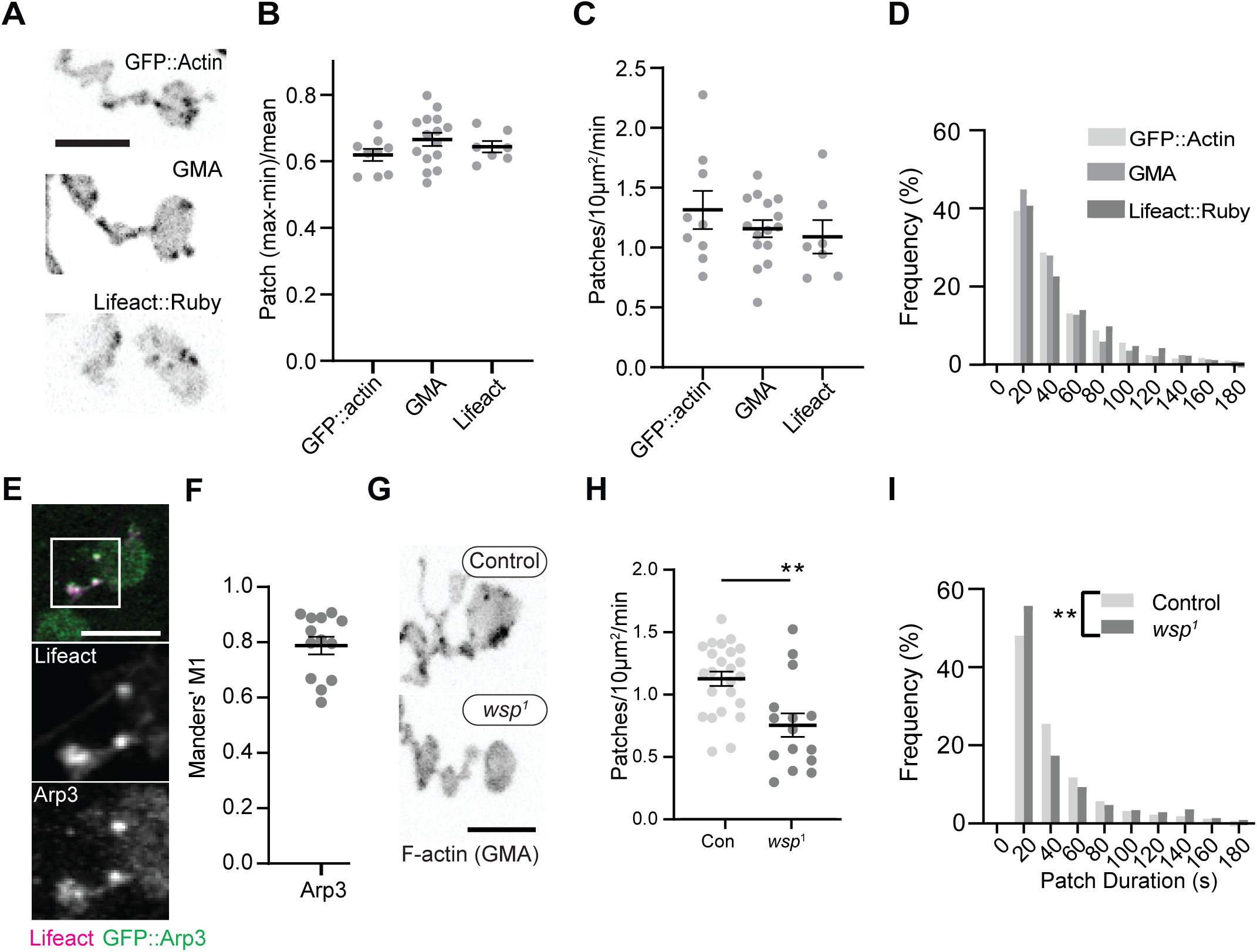
Dynamics of synaptic actin structures are consistent with endocytic function. (A) Representative MaxIPs of single spinning disk confocal microscopy time points, showing C155-Gal4-driven actin probes: GFP::actin, GMA, and Lifeact::Ruby. (B-D) Automatic detection and analysis of movies acquired at .25Hz of F-actin patch intensity amplitude (B), frequency (C), and duration distribution (D) show similar dynamics for different reporters. (E-F) MaxIPs (Airyscan microscopy) of a single frame of muscle 6/7 NMJ expressing Lifeact::ruby (magenta) and Arp3::GFP (green). Actin patches colocalize extensively with Arp3::GFP. (F) Quantification of colocalization by Manders’ coefficients. Graph shows mean +/− s.e.m.; n represents NMJs. (G-I) Patch assembly requires the Arp2/3 activator WASp. GMA patch dynamics in control and WASp mutant animals imaged at .25Hz. (G) MaxIPs of single spinning disk confocal microscopy time points, showing pan-neuronally expressed GMA localization in control and *wsp*^1^ mutant muscle 6/7 NMJs. (H) Quantification of patch frequency. Graph shows mean +/− s.e.m.; n represents NMJs. (I) Quantification of patch duration distribution. Bins are 20 sec; X axis values represent bin centers. n represents patches. Scale bars in A, E, G, are 5 μm. Associated with Fig. 1 S1 and Movie 1.

We next examined the molecular determinants of synaptic actin patch assembly. Patches strongly co-labeled with Arp3::GFP (Manders’ coefficient of 0.81 (Lifeact:Arp3), (**Fig 1E-F**)), suggesting they are predominantly composed of branched F-actin, similar to sites of endocytosis in other cell types (Akamatsu et al., 2020; Collins et al., 2011). To test whether synaptic actin patches require Arp2/3 activation, we analyzed patch dynamics in larvae lacking the Arp2/3 activator WASp. We compared a genomic mutant (**Fig 1G-I)**, likely hypomorphic due to maternal contribution (Ben-Yaacov et al., 2001)) to presynaptic depletion in neurons expressing WASp RNAi (**Fig 1 S1C-E**). Using both approaches allows us to distinguish neuron-autonomous from non-autonomous effects of WASP, which is present both pre- and postsynaptically (Coyle et al., 2004). Both genomic and RNAi manipulations significantly reduced the number of actin patches, while genomic mutants also skewed the distribution of patch durations towards both shorter and longer events (**Fig 1I**). These differences could reflect variable loss of function between the RNAi and mutant, or identify separable presynaptic autonomous (patch frequency), vs non-autonomous (patch duration) effects of WASp. Overall, these data clearly indicate that WASp is autonomously required in neurons to initiate assembly of presynaptic actin patches, similar to its involvement in endocytosis in yeast, mammalian non-neuronal cells, and in the NMJ (Hussain et al., 2001; Kessels and Qualmann, 2004; Khuong et al., 2010; Madania et al., 1999).

We next examined the synaptic distribution of two likely WASp regulators, Nwk and Dap160. By conventional and super-resolution microscopy of neurons in diverse organisms, these and other presynaptic membrane remodeling proteins localize to a broad membrane domain surrounding active zones, termed the periactive zone (PAZ) (Coyle et al., 2004; Denker et al., 2011; Gerth et al., 2017; Koh et al., 2004; Marie et al., 2004; Sone et al., 2000). Consistent with these prior descriptions, we observed by structured illumination microscopy (SIM) that the PAZ proteins Nwk and Dap160 localize to a membrane-proximal mesh that surrounds active zones, which were labeled with Bruchpilot (BRP, **Fig 2A**). We observed similar results by live imaging of an endogenously tagged Nwk protein by SIM, which revealed the vast majority of protein to be in close proximity to the plasma membrane (**Fig 2B**). We then compared the localization of PAZ proteins to F-actin patches at the NMJ. As expected, actin patches were much sparser than the endocytic machinery, and GMA-labeled patches only partially overlapped with each of presynaptic WASp, Nwk, and Dap160 (**Fig 2C-F)**; Manders’ coefficients of 0.45, 0.59, 0.53, respectively). These data confirm that, in sharp contrast to the actin regulatory machinery, which localizes broadly across the PAZ, actin assembly itself is much sparser both spatially and temporally at the NMJ. This raises the question of how PAZ machinery might itself be locally regulated to promote the formation of productive synaptic actin assemblies.

**Figure 2.**
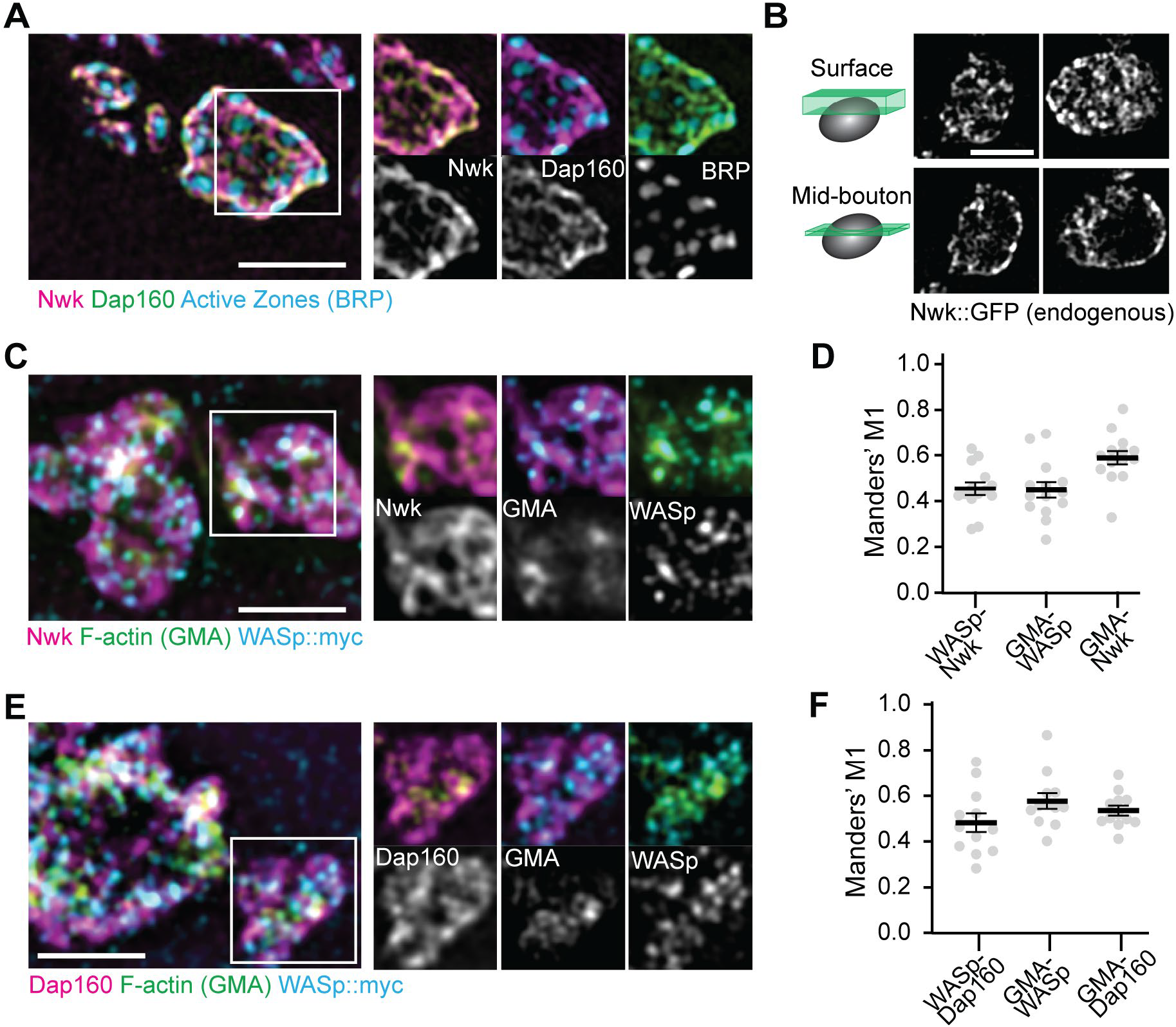
Periactive zone proteins accumulate broadly across the NMJ. (A-B) The periactive zone (PAZ) proteins Nwk (magenta) and Dap160 (green) accumulate in a micron-scale mesh adjacent to active zones (AZ) (Bruchpilot, blue). Image shows maximum intensity projection (MaxIP) of a structured illumination microscopy (SIM) Z-stack. (B) Surface projection (top) and medial optical section (bottom) SIM images of live-imaged endogenous Nwk::GFP showing abundant and specific membrane recruitment, similar to fixed imaging. (C-F) PAZ proteins partially colocalize with actin patches. Optical slices of SIM micrographs showing F-actin (labeled with GMA) localization with presynaptically expressed WASp::Myc and Nwk (C) or Dap160 (E). (D,F) Quantification of colocalization between GMA and WASp::Myc, and Nwk (D), or Dap160 (F). (D,F) Graphs show mean +/− s.e.m.; n represents NMJs. Associated with **Fig 1 S1**.

### Multiple interaction interfaces between Dap160 and Nwk regulate Nwk autoinhibition

The hypothesis that PAZ protein-mediated actin assembly might be locally activated is particularly interesting given that we and others have previously shown that autoinhibition of both Nwk and its mammalian homolog FCHSD2 suppresses both WASp activation and membrane binding (See **Fig 3A** for summary model; Almeida-Souza et al., 2018; Rodal et al., 2008; Stanishneva-Konovalova et al., 2016). These results suggest that transient or localized relief of autoinhibition could explain how the PAZ controls actin assembly. To determine if and how the candidate activator Dap160 might relieve Nwk autoinhibition, we first mapped their specific interaction domains using GST pulldown assays, and found that purified Dap160 SH3C-containing protein fragments (SH3C, SH3CD, or SH3ABCD) directly interact with Nwk^SH3b^, while SH3D alone does not (**Fig 3B, Fig 3S1A-C**, see **Fig 3 S2A** for details of constructs used). Unexpectedly, Dap160 SH3C, SH3D, and SH3CD domain fragments also each interact with the isolated Nwk F-BAR domain (**Fig 3S1B**). We next determined how Dap160 interactions with Nwk^F-BAR^ compared to a Nwk fragment containing the F-BAR and both SH3 domains. Dap160-Nwk^F-BAR^ interactions were progressively eliminated by increasing salt, suggesting they are mediated by electrostatic interactions. By contrast, Dap160^SH3CD^-Nwk interactions were maintained (**Fig 3B, Fig 3S1C**), suggesting that the SH3-SH3 interaction is mediated primarily by hydrophobic interactions, consistent with their mammalian homologs (Almeida-Souza et al., 2018) (see summary of interactions, **Fig 3C**). Finally, we found that truncation of Dap160^SH3CD^ significantly decreased the levels of Nwk in synaptic boutons (**Fig 3D, Fig 3S2**) as well as colocalization between Nwk and Dap160 (**Fig 3E**), supporting an *in vivo* requirement for this interaction.

**Figure 3.**
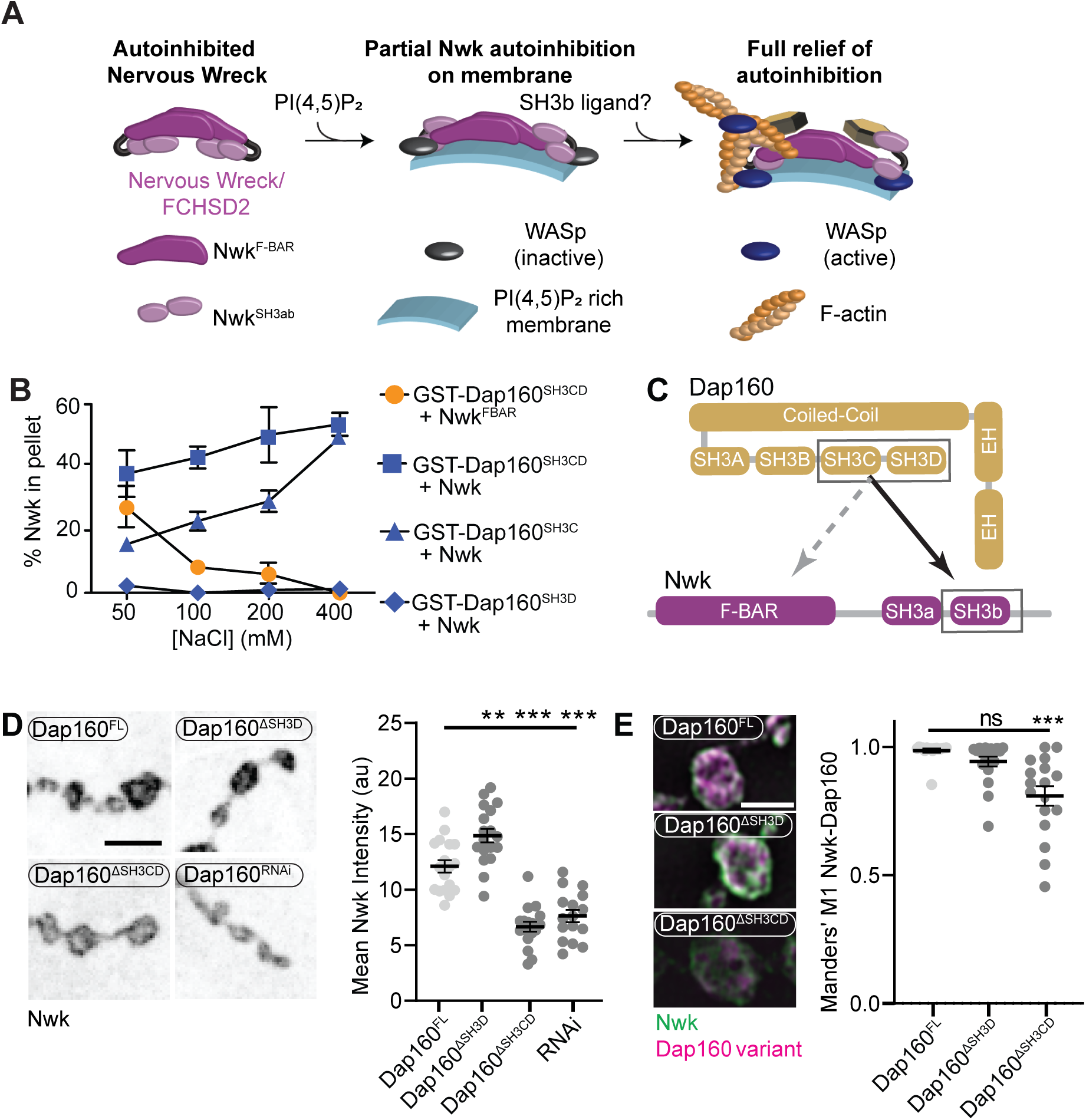
Distinct SH3-SH3 and SH3-BAR domain interactions drive Dap160-Nwk association *in vitro* and at synapses. (A) Model for autoinhibition of Nwk membrane binding and WASp activation. Neither membrane-bound nor membrane-free Nwk efficiently activate WASp-mediated actin polymerization, due to persistent SH3b-mediated autoinhibitory interactions, suggesting that an SH3b domain ligand is required for activation. (B) Dap160^SH3CD^ exhibits electrostatic and hydrophobic interactions with the Nwk F-BAR and SH3 domains, respectively. GST fusion proteins were immobilized on glutathione agarose and incubated with the indicated purified proteins. Pellets and supernatants were fractionated by SDS-PAGE, Coomassie stained, and quantified by densitometry. Graphs show the average +/− s.e.m. of three independent reactions. [Nwk^F-BAR^]=1.5 μM, [Nwk]=.8μM, [GST-Dap160^SH3CD^]=1.6μM, [GST-Dap160^SH3C/D^]=1.2μM. (C) Summary of Dap160^SH3CD^-Nwk^SH3ab^ interactions. Gray and black arrows indicate electrostatic and hydrophobic interactions, respectively (D) MaxIP spinning disc confocal or (E) MaxIP SIM micrographs of muscle 4 NMJs expressing C155-GAL4-driven UAS-Dap160 rescue transgene variants in a *dap160* null background (dap160^Δ1/Df^). Loss of the Dap160^SH3CD^ domains (Dap160^ΔSH3CD^), but not the SH3D domain alone (Dap160^ΔSH3D^), disrupts normal Nwk accumulation (D, right) and Dap160-Nwk colocalization (E, right) at synapses. Graphs show mean +/− s.e.m.; n represents NMJs. Associated with **Fig S3S1-2**.

### Dap160 ^SH3CD^ and membranes relieve inhibition of Nwk-WASp-Arp2/3 actin assembly *in vitro*

We previously showed that Nwk only weakly activates WASp-dependent actin assembly *in vitro*, due to Nwk autoinhibition (Stanishneva-Konovalova et al., 2016). To test whether Dap160^SH3CD^ might relieve Nwk autoinhibition, we performed pyrene-actin assembly assays (**Fig 4**). At moderate Nwk-Dap160 concentrations (500nM and 2um, respectively), Nwk and Dap160^SH3CD^ significantly enhanced the rate of WASp-Arp2/3-mediated actin assembly compared to Nwk plus WASp alone (**Fig 4A**). This effect is through Nwk, as Dap160^SH3CD^ had no effect on WASp-Arp2/3 in the absence of Nwk. Further, Dap160 enhancement of Nwk-WASp actin assembly required the Dap160^SH3D^ domain, further showing that the specific Dap160^SH3D^-Nwk^F-BAR^ interaction relieves functional Nwk autoinhibition *in vitro*. Thus, multiple Nwk-Dap160 interactions work together to relieve autoinhibition of Nwk.

**Figure 4.**
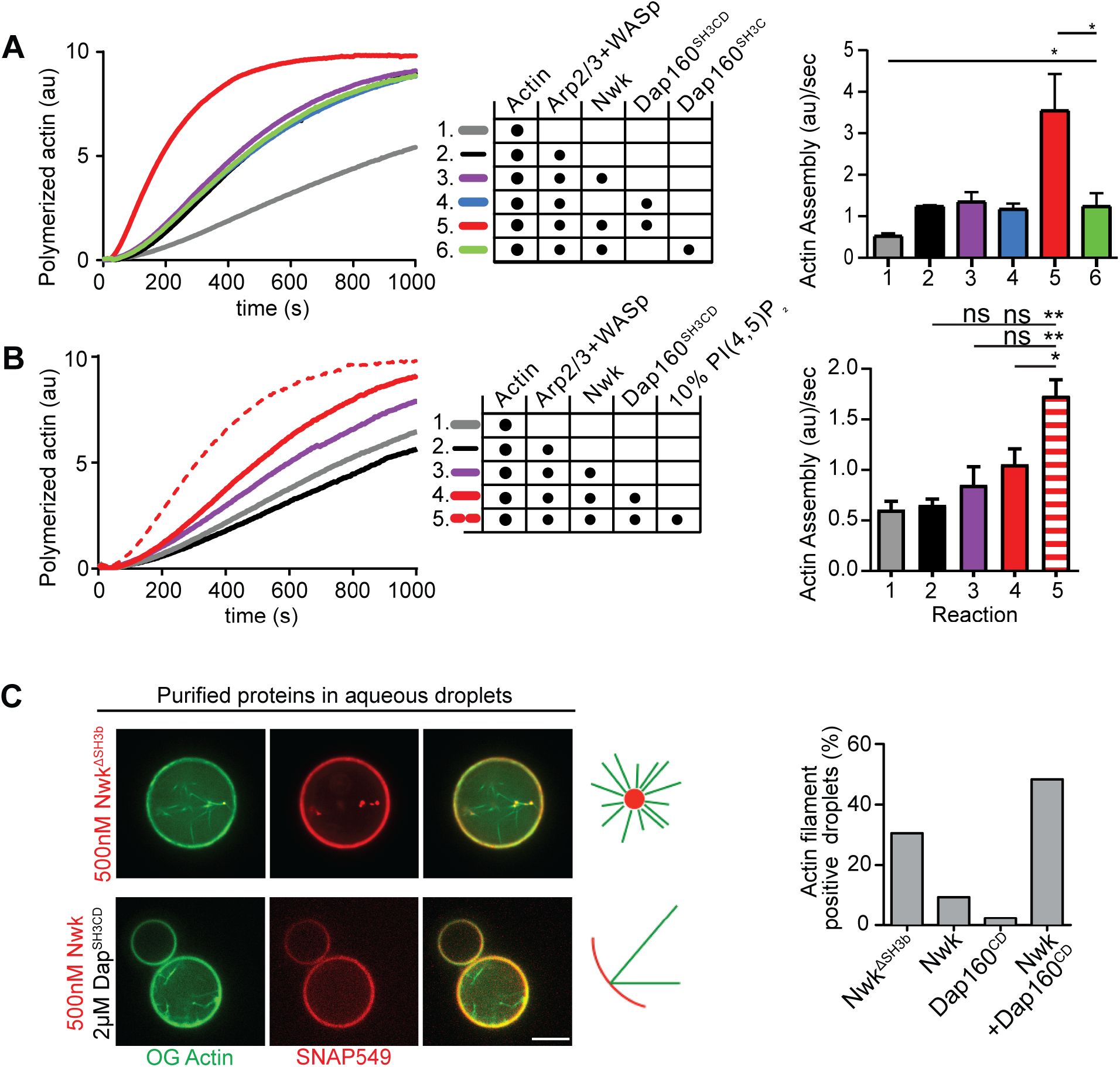
Nwk, Dap160, and PI(4,5)P_2_ potentiate WASp-mediated actin assembly at membranes. (A-B) Pyrene actin assembly assay (2.5 μM actin, 5% pyrene labeled). Curves are representative single experiments demonstrating actin assembly kinetics; graphs represent rates calculated from the linear range of assembly from at least two independent experiments. (A) The combination (red trace) of Nwk and Dap160^SH3CD^ together enhance WASp-Arp2/3 mediated actin assembly. Either alone (magenta and blue traces) has no effect on WASp activity. [Nwk] = 500nM, [Dap160] = 2 μM, [WASp] = 50 nM, [Arp2/3] = 50 nM. (B) PI(4,5)P_2_ enhances Nwk-Dap160 activation of WASp-mediated actin assembly. Nwk alone or in combination with 10% PI(4,5)P_2_ liposomes fails to activate WASp, while the addition of Dap160^SH3CD^ and PI(4,5)P_2_ synergistically enhance WASp-mediated actin assembly. [Nwk] = 100 nM, [Dap160] = 500 nM, [WASp] = 50nM, [Arp2/3] = 50 nM. (C) Single slices from spinning disc confocal micrographs of water-droplet actin assembly assay: SNAP labeled Nwk constructs (red) and Oregon Green actin (green) were mixed with the indicated proteins in aqueous solution and emulsified in 97.5% DPHPC, 2.5% PI(4,5)P_2_ in decane. Both deregulated Nwk^ΔSH3b^ and Nwk + Dap160^SH3CD^ promote F-actin assembly in droplets. However, while Nwk-Dap160^SH3CD^ derived F-actin associates with the lipid interface, de-regulated Nwk^ΔSH3b^ promotes actin assembly from asters that do not associate with membrane. [Nwk^1-xxx^] = 500nM, [Dap160] = 2μM, [WASp] = 50nM, [Arp2/3] = 50nM. Graph indicates percentage of droplets with observable actin filament assembly. Scale bar in C is 10 μm.

To generate salient physiological force, actin assembly must be coupled to membranes, and negatively charged lipids are an important ligand for both Nwk and WASp. Thus, we next tested whether addition of PI(4,5)P_2_-rich liposomes modified actin assembly by Nwk, Dap160, and WASp (**Fig 4B**). Indeed, PI(4,5)P_2_ containing liposomes synergistically activated WASp-mediated actin assembly in concert with Dap160 and Nwk. By contrast, neither Nwk, PI(4,5)P_2_, or Nwk+PI(4,5)P_2_ on their own were sufficient to activate WASp above baseline (**Fig 4B**). Since PI(4,5)P_2_ is also insufficient to robustly activate either WASp or Nwk under these conditions (Stanishneva-Konovalova et al., 2016), our data suggest that WASp activation reflects coordinated relief of Nwk autoinhibition by both Dap160 and membranes. To further explore the coupling between lipid association and actin assembly, we conducted F-actin assembly assays in a droplet assay, in which protein-containing aqueous droplets are surrounded by a lipid interface, with lipid head groups directly contacting the aqueous phase (**Fig 4C**). In this assay, we found that coordinated interactions among Nwk, Dap160, and WASp directed actin assembly to the lipid interface. By contrast, substitution of Nwk lacking its autoinhibitory and Dap160-interacting SH3b domain (Nwk^ΔSH3b^) caused actin to assemble as free-floating asters (**Fig 4C**). We previously found that expression of a similarly deregulated fragment (Nwk^1-631^) at the NMJ led to diffuse actin filament assembly throughout the synapse (Stanishneva-Konovalova et al., 2016). Together these data suggest that Nwk^SH3b^ has a dual role of maintaining autoinhibition via Nwk-F-BAR interactions and permitting actin assembly at specific synaptic locations via Dap160-mediated activation.

### Dap160 and WASp relieve Nwk autoinhibition and promotes its membrane association

Our actin assembly data suggest that membrane recruitment is a critical regulator of the Nwk-Dap160-WASp complex (**Fig 4B-C**). To test whether Nwk-Dap160 interactions directly regulate membrane recruitment, we performed liposome co-sedimentation assays. We found that Dap160^SH3CD^ enhanced Nwk membrane binding in a dose-dependent fashion (**Fig 5A**). This effect depended on membrane charge, as Dap160^SH3CD^ significantly enhanced Nwk membrane binding at both 5% and 10%, but not at 2.5% PI(4,5)P_2_ (**Fig 5B**). Only at 10% PI(4,5)P_2_ did Dap160^SH3CD^ promote Nwk membrane binding to the same extent as the completely uninhibited Nwk^FBAR^ domain alone, suggesting that membrane charge and intermolecular interactions with Dap160 together tune Nwk membrane recruitment. Indeed, this effect required the full Dap160^SH3CD^-Nwk^SH3b^ interaction: Dap160^SH3C^ alone was unable to promote membrane binding by Nwk, and Dap160^SH3CD^ did not enhance membrane binding of Nwk lacking its Dap160-interacting SH3b domain (**Fig 5S1A**). These data further support the hypothesis that Dap160^SH3CD^ relieves Nwk^SH3b^ mediated autoinhibition.

**Figure 5.**
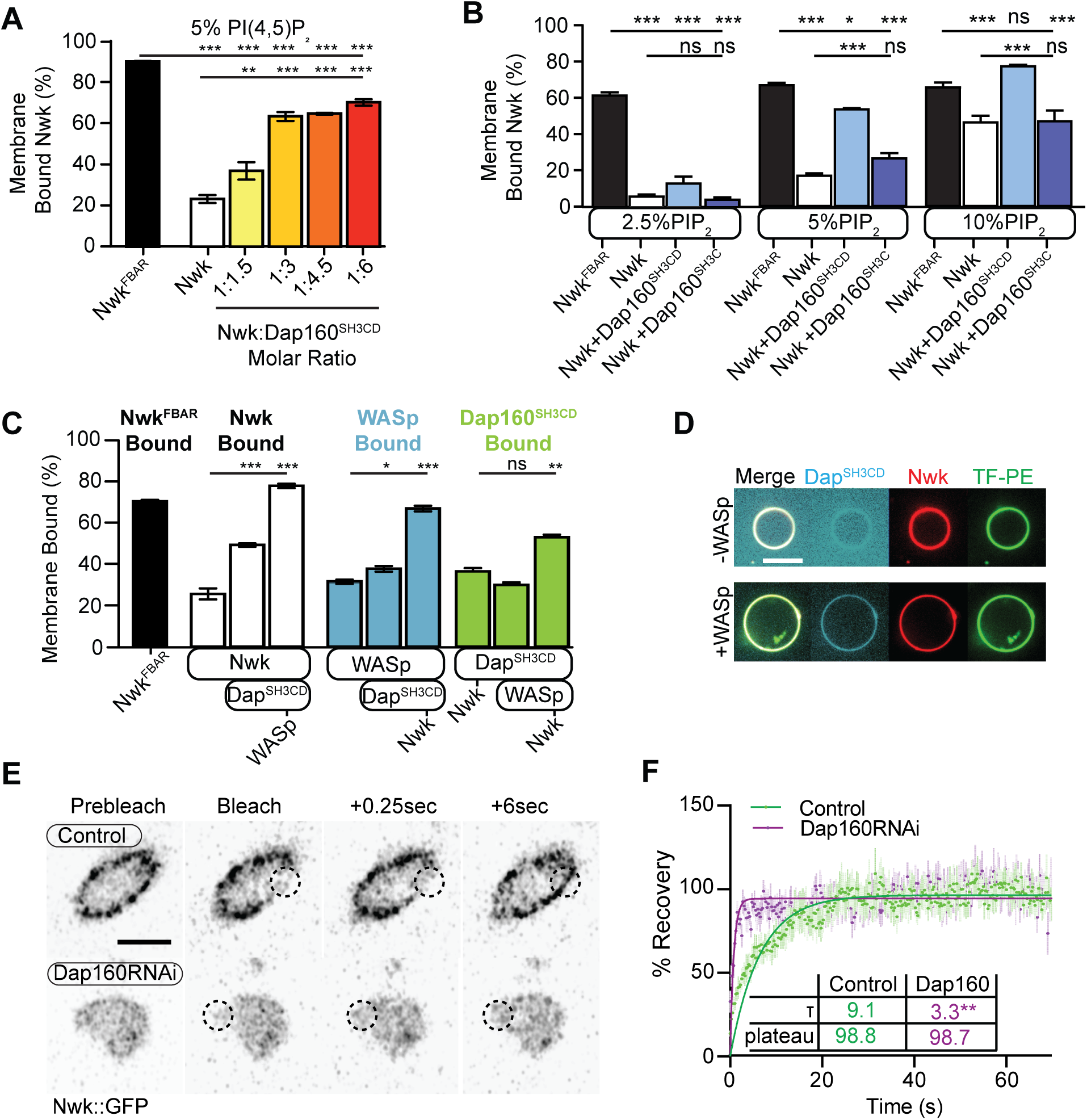
Dap160^SH3CD^ and WASp promote Nwk membrane association. (A-C) Liposome co-sedimentation assays between the indicated purified proteins and liposomes composed of [mol% = DOPC/DOPE/DOPS/PI(4,5)P_2_ = 80-x/15/5/x], with x representing PI(4,5)P_2_ concentration as noted. Quantification from Coomassie-stained gels represents the mean fraction of total protein that cosedimented with the liposome pellet, ± SEM. (A) 1:3 Nwk:Dap160^SH3CD^ saturates enhancement of Nwk membrane association at 5% PI(4,5)P_2_, but not to the level of the isolated Nwk F-BAR alone (Nwk^F-^ BAR, black bar). [Nwk^1-xxx^] = 2 μM, [Dap160] = 6 μM. (B) Dap160^SH3CD^ (but not Dap160^SH3C^) enhances Nwk association with membranes at a range of PI(4,5)P_2_ concentrations. Maximum binding (comparable to Nwk^F-BAR^ occurs only at 5-10% PI(4,5)P_2_ concentrations. [Nwk^F-BAR^] = 3 μM, [Nwk] = 1.125 μM, [Dap160^SH3CD^] = 1.7-6.8 μM. (C) Nwk, WASp, and Dap160^SH3CD^ mutually enhance membrane recruitment. Addition of Dap160^SH3CD^ and WASp additively enhance Nwk membrane association, while Dap160^SH3CD^ and WASp show maximum recruitment to 10% PI(4,5)P_2_ liposomes in the presence of both other proteins. [Nwk] = 1 μM, [WASp] = 1 μM, [Dap160^SH3CD^] = 3 μM. (D) GUV decoration assay, with 10% PI(4,5)P_2_ GUVs labeled with <1% TopFluor-PE. The addition of WASp to Nwk (red) and Dap160^SH3CD^ (blue) enhances the recruitment of Dap160^SH3CD^ to the membrane (green, note diffuse blue signal in (-) WASp condition). [Nwk] = 250 nM, [WASp] = 250 nM, [Dap160^SH3CD^] = 1 μM Scale bar is 10 μm (E-F) FRAP assay of endogenously labeled Nwk in control and C155-GAL4/UAS-Dicer driven Dap160^RNAi^ NMJs. Images show individual medial optical sections of Airyscan confocal images at the indicated time point. Control Nwk signal shows strong membrane association (see strong peripheral signal) and slower recovery kinetics, while loss of Dap160 eliminates the strong peripheral accumulation of Nwk::GFP and increases the recovery kinetics of Nwk::GFP in the bleached region (dashed circles). Graph shows mean +/− s.e.m.; n represents NMJs. Scale bar is 5 μm.

As we found that Dap160^SH3CD^ is insufficient to fully activate membrane binding by Nwk at intermediate phosphoinositide concentrations (**Fig 5A**), we asked whether WASp could further enhance Nwk membrane recruitment. Indeed, the addition of Dap160^SH3CD^ and WASp together enhanced Nwk membrane association to the level of the isolated F-BAR domain (**Fig 5C**). Moreover, coordinated binding of all three components resulted in significantly enriched membrane association of both WASp and Dap160 (**Fig 5C**). We directly observed the coordinated recruitment of Nwk and Dap160 in the presence of WASp using fluorescently labeled proteins on GUVs (**Fig 5D**). Consistent with the direct Dap160-Nwk^SH3b^ interaction, we found that deletion of the Nwk^SH3b^ domain abolished both the Dap160^SH3CD^-dependent increase and the coordinated recruitment of WASp and Dap160 (**Fig 5S1A**). Notably, addition of Dap160 and WASp did not change the nature of membrane deformations generated by Nwk (scalloped and pinched membranes (Becalska et al., 2013)), suggesting that Dap160 and WASp together potentiate rather than alter the inherent activity of Nwk (**Fig 5S1D**). These data indicate that Dap160-Nwk SH3-mediated interactions potentiate Nwk association with membranes *in vitro*.

Finally, to test whether Dap160 promotes Nwk membrane association *in vivo*, we examined the dynamics of Nwk at the synapse in the presence and absence of Dap160. Knockdown of Dap160 by RNAi (**Fig 5E**) led to a striking loss of endogenously tagged Nwk::GFP from synaptic membranes (note strong peripheral labeling in control bouton cross-sections, **Fig 5E**). Further, Dap160 knockdown significantly increased the rate of recovery of Nwk::GFP, consistent with a shift in localization from membrane-bound to cytosolic (**Fig 5F**). These data suggest that the Dap160^SH3CD^-Nwk interaction promotes Nwk membrane association *in vivo*.

Taken together, our data indicate that multiple coordinated interactions between Nwk, WASp, Dap160^SH3CD^, and membranes are required to relieve Nwk autoinhibition, allowing for tight control of membrane-coupled actin assembly in the PAZ.

### Dap160-Nwk interactions regulate synaptic F-actin patch dynamics

To determine how these mechanisms direct WASp-mediated actin assembly at the synapse, we measured actin dynamics in *nwk* (**Fig 6A-C, Movie 2**) and *dap160* domain (**Fig 6D-F**) mutant NMJs. We predicted two possible but non-exclusive functions based on the dual roles that we found for the Nwk-Dap160-WASp module *in vitro*: If Nwk and Dap160 are primary activators of WASp, then loss of function mutants are likely to diminish patch frequency, duration, or intensity. Importantly, multiple WASp activators exist in the synaptic endocytic machinery (e.g. Cip4 and Snx9 (Almeida-Souza et al., 2018; Gallop et al., 2013)), and therefore it would be possible that these could make significant contributions to WASp activation in addition to Nwk. Conversely, if an important function of autoinhibition is to ‘clamp’ actin assembly at the synapse, we expect that loss of Nwk and/or Dap160 would lead to spurious actin assembly events by these other WASp regulators. Strikingly, both *nwk* and Dap160^ΔSH3CD^ mutants significantly increased patch frequency (**Fig 6B,E**), supporting a clamp function for these proteins. We did not find a difference in the distribution of patch lifetimes, suggesting that it is the frequency of events, and not their duration *per se*, that changes (**Fig 6C,F**).

**Figure 6.**
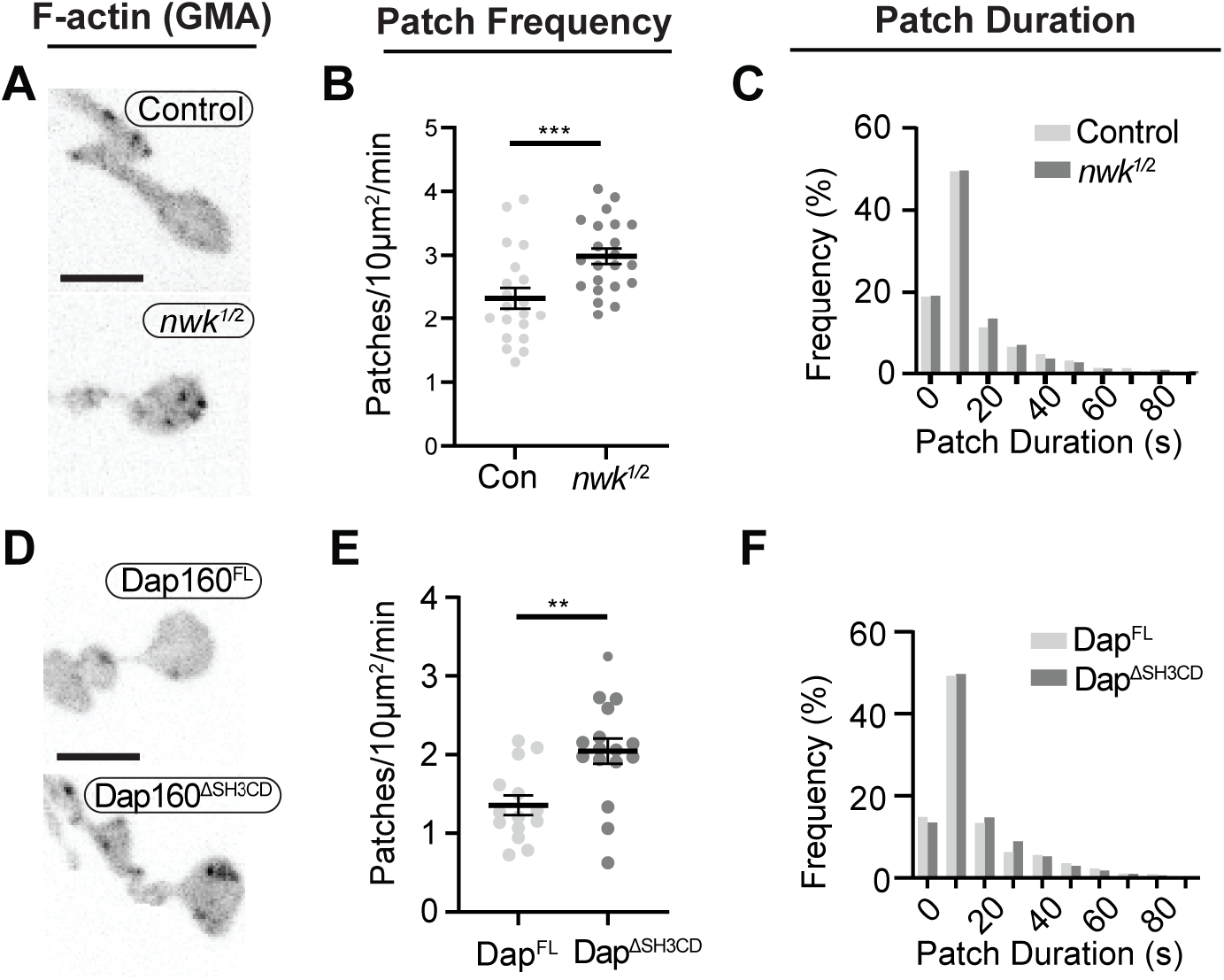
Loss of the Dap160-Nwk interaction disrupts actin patch dynamics at synapses *in vivo*. Quantification of GMA-labeled F-actin patch dynamics. (A,C) MaxIPs of live spinning disc confocal micrographs of GMA in the indicated genotypes, imaged at 1Hz. Graphs quantify patch frequency (B, E) and distribution of patch durations (C, F) and. Loss of each of *nwk* (A-C) increases the frequency of patches. Similarly, loss of the Nwk-interacting Dap160^SH3CD^ domain increases the frequency of patch assembly events, with no change in patch durations. Scale bars in (A,D) are 5 μm. Associated with **Fig 6 S1**, **Movie 2**.

We also analyzed actin dynamics using a complementary approach in which we measured the normalized intensity variation (CoV) over time across the entire NMJ. Interestingly, the magnitude of variation was significantly higher in *nwk* mutants (**Fig 6S1A-B**), but the area of the NMJ that was highly variant was similar between genotypes, suggesting that actin assembly is more dynamic in time in these mutants, rather than more extensive in space (**Fig 6S1C**). We validated this analysis for its sensitivity in detecting changes in event frequency by analyzing synthetic data (**Fig 6S1D**, see Methods for detail). The modeled data suggest that the difference in CoV that we measure between Control and nwk is consistent with a 43% increase in patch frequency, which is slightly higher than our measurement by particle tracking (28%, **Fig 6A**). This complementary analysis does not rely on particle tracking and makes no assumptions about the nature of actin dynamics, and is consistent with our particle-based metrics. Thus, we conclude that these phenotypes are robust to the method of analysis used.

### Nwk and Dap160 ^SH3CD^ are required for normal synaptic vesicle endocytosis

We next investigated the physiological function of actin patches *in vivo*. Considering that patch morphology, frequency, and duration resembled endocytic dynamics, we first compared actin patches with the endocytic adapter AP2α. Like other endocytic proteins, endogenously tagged AP2α::GFP was highly abundant at the synapse and localized across much of the presynaptic membrane both diffusely and in highly dynamic puncta, only a subset of which dynamically colocalized with actin patches (**Fig 7A-B, Movie 3**). By contrast, a majority of actin patches colocalized with AP2α::GFP (**Fig 7B-C**, Manders’ M1 = 0.59+/− .09) in single live images, consistent with a role in endocytosis for these actin-enriched sites. To functionally test the hypothesis that actin patches are endocytic, we acutely disrupted endocytic dynamics using the temperature sensitive dominant negative dynamin*/shi*^TS1^ allele. When imaged under restrictive conditions, *shi* disruption decreased the frequency of actin patch dynamics (**Fig 7D-E**). Together, these data suggest that a significant fraction of presynaptic actin patches are associated with endocytosis.

**Figure 7.**
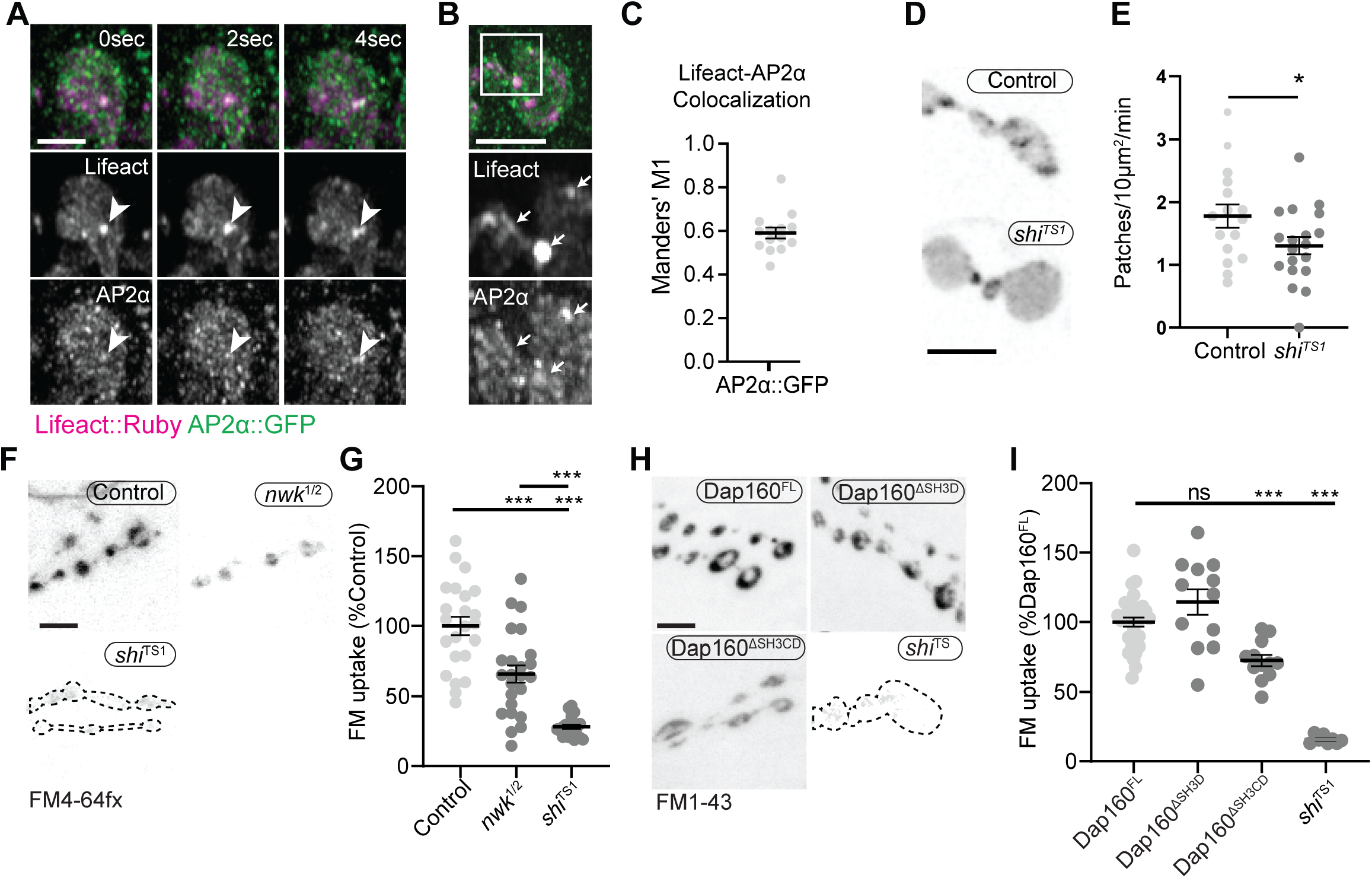
Actin patches and the Nwk-Dap160 interaction are associated with synaptic endocytosis. (A-B) MaxIPs of Airyscan micrographs of endogenously tagged AP2α::GFP (green) and Lifeact::Ruby (magenta) in live (A) or fixed (B) preparations. Note AP2α::GFP knockin is endogenously expressed in both neurons and muscle. (A) Timelapse showing dynamic colocalization of Lifeact::Ruby actin patches with AP2α::GFP. (B-C) A majority of actin patches colocalize with presynaptic AP2α::GFP signal. (C) Quantification of colocalization by Manders’ coefficients. M1 represents % of lifeact region that contains AP2α::GFP signal above threshold. (D-E) Normal patch assembly requires Dynamin activity. (D) MaxIPs of single spinning disk confocal microscopy time points, showing pan-neuronally expressed GFP::actin in control and *shi*^TS1^ mutant m6/7 NMJs, imaged at 1 Hz, at the restrictive temperature of 31° under stimulating conditions to drive the terminal *shi*^TS1^ phenotype (45mM KCl, 2mM CaCl2). Graph shows mean +/− s.e.m. n represents NMJs. (E) Quantification of patch frequency. (F-I) FM dye uptake assays at m6/7 NMJs following 5 min 90 mM potassium stimulation at 36°C. (A,C) MaxIPs of spinning disc confocal micrographs of FM dye uptake assays. (A-B) *nwk* mutants exhibit partially defective FM4-64fx dye uptake relative to *shi*^TS1^ mutants. (C-D) Loss of Dap160-Nwk interactions in a Dap160^ΔSH3CD^ truncation (but not Dap160^ΔSH3D^) exhibit partially defective FM1-43 dye uptake relative to *shi*^TS1^, similar to *nwk* mutants. Graphs show mean +/− s.e.m.; n represents NMJs. Scale bars are 2.5 μm (A, B) or 5 μm (D, F, H). Associated with **Fig 7 S1-2**.

We next tested the physiological requirement of the Nwk and Dap160^SH3CD^ interaction. As both Nwk and Dap160 are implicated in the endocytic trafficking of synaptic growth-promoting BMP receptors (O’Connor-Giles 2008, Rodal 2008), we tested whether the Dap160-Nwk interaction was required for normal synaptic growth, which we assayed by counting satellite boutons, a hallmark phenotype of both null mutants. Surprisingly, we found that both Dap160^ΔSH3D^ and Dap160^ΔSH3CD^ truncations rescued satellite bouton numbers to wild type levels (**Fig 7S2**), suggesting that actin dynamics phenotypes in the Dap160^ΔSH3CD^ mutant are not associated with synaptic growth regulation. We next examined synaptic vesicle endocytosis and recycling by FM dye uptake. *nwk*^1/2^ null mutants led to a 34% decrease in FM4-64fx uptake compared to controls (**Fig 7A-B**), an intermediate phenotype compared to dominant negative dynamin in *shi*^TS1^ mutants (FM uptake 28% of controls). *dap160* null mutants have been previously shown to exhibit an endocytosis defect (Koh et al., 2004; Marie et al., 2004), so we next tested whether the interaction between Dap160 and Nwk is required to support normal endocytosis. Indeed, we found that expression Dap160^ΔSH3CD^ in *dap160* null mutants also significantly diminished FM dye uptake to a similar extent as loss of *nwk* (27% reduction, **Fig 7C-D**). By contrast, loss of the Dap160^SH3D^ domain alone caused no defects in FM uptake, consistent with the lack of effect of this mutation on Nwk accumulation and localization (**Fig 3D-E**), and suggesting that this interaction, though required *in vitro*, may be compensated by additional factors *in vivo*. Both *nwk* and Dap160^ΔSH3CD^ mutants unloaded FM dye to the same extent as controls, suggesting that diminished endocytosis is a direct phenotype, and not secondary to exocytic deficits (**Fig 7S1**). Importantly, these data indicate that spurious actin assembly events in *nwk* and *dap160* mutants are likely to be unproductive for normal endocytosis. Overall, our data support the hypothesis that normal synaptic actin patches represent active endocytic events, and indicate that Dap160-Nwk regulation of actin patch dynamics is functionally required for synaptic vesicle endocytosis.

## Discussion

Here we identify a mechanism by which autoinhibition clamps the presynaptic endocytic machinery to regulate the dynamics of discrete synaptic actin assembly events and the efficiency of synaptic endocytosis. We identify specific interactions among Nwk, Dap160, and WASp that function in two ways to potentiate membrane associated actin dynamics: 1) Persistent autoinhibition of Nwk allows for stable binding of inactive PAZ machinery to presynaptic membranes to constrain spurious actin assembly events. 2) Coordinated relief of Nwk autoinhibition by Dap160 and WASp robustly activate F-actin assembly and ensure that actin assembles into structures that are likely to productively remodel membranes. This provides a mechanism by which synapses can use the micron-scale PAZ organization of endocytic machinery as a regulated reservoir to efficiently generate 50-100 nm-scale endocytic events, in response to physiological cues such as synaptic transmission.

### The predominant presynaptic actin structures resemble endocytic patches

Here we provide the first quantitative analysis of the composition and dynamics of individual presynaptic F-actin structures. Numerous studies have examined actin dynamics at the level of entire synapses or qualitatively described dynamics of discrete actin structures (Bloom et al., 2003; Colicos et al., 2001; Nunes et al., 2006; Piccioli and Littleton, 2014; Sankaranarayanan et al., 2003; Zhao et al., 2013), and identified diverse roles for actin, including synaptic vesicle endocytosis (Holt et al., 2003; Kononenko et al., 2014; Soykan et al., 2017; Watanabe et al., 2013; Wu et al., 2016; Zhao et al, 2013), synaptic vesicle organization and mobilization (Guzman et al., 2019; Lee et al., 2012; Marra et al., 2012; Owe et al., 2009; Sakaba and Neher, 2003; Wolf et al., 2015), active zone organization and function (Boyl et al., 2007; Morales et al., 2000; Wagh et al., 2015; Waites et al., 2011; Wang et al., 1999), and receptor-mediated endocytosis (Kim et al., 2019; Rodal et al., 2008). Bulk analyses, which do not separate individual dynamic actin structures in space and time, are limited in their ability to discern how the regulation and dynamics of actin contribute to these distinct functions. We leveraged our ability to extract data describing individual structures to find that synaptic actin predominantly assembled into discrete Arp2/3-associated patches, and identified points of control over their dynamics. Specifically, we found that loss of endocytic proteins differentially affected the frequency and kinetics of individual actin patches, which correlate with functional deficits in endocytosis.

The link between the actin structures that we observe and endocytic events is supported by several lines of evidence: The morphology and duration of synaptic actin patches are similar to WASp/Arp2/3-dependent endocytic actin dynamics in cultured mammalian cells (~40 seconds; Merrifield et al., 2004; Taylor et al., 2011). The frequency of patch assembly, which we measured in resting synapses, approaches the rate of spontaneous synaptic vesicle release at this synapse (**Fig S1G**) (~5-6/10 μm^2^/min; Melom et al., 2013, Akbergenova 2018). Further, actin patches colocalize partially with endocytic adapters and their assembly is sensitive to disruption of endocytosis (**Fig 7D-E**). Finally, we found that the same endocytic proteins and protein interactions that regulate endocytosis at this synapse also alter the dynamics of actin patches.

Technical challenges due to the high density of endocytic proteins and synaptic vesicle cargoes, and the difficulty of conducting sparse single vesicle measurements at this synapse (compared to neurons in culture (Chanaday and Kavalali, 2018; Peng et al., 2012)) make it difficult to directly link the dynamics of actin structures to specific membrane or cargo internalization events. However, the frequency of the events captured by our approach makes it unlikely that they represent rare F-actin-dependent events at this synapse, such as those that control macropinocytosis or new bouton growth (Khuong et al., 2010; Kim et al., 2019; Piccioli and Littleton, 2014), and more likely that they represent bona fide endocytic events. Thus, while we do not rule out other biological functions for a subset of patches, together our data indicate that a significant and measurable fraction of synaptic actin patches are associated with endocytosis.

### Autoinhibition clamps PAZ membrane remodeling machinery at synapses

Our data demonstrate that much of the synaptic membrane remodeling machinery is held in an inactive state at the presynaptic membrane: Nwk and Dap160 accumulate in a micron-scale membrane domain (**Figs 1A, S1A, 3C**), and loss of this machinery increases the frequency of short-lived actin patches (**Fig 6**). These data suggest that these PAZ proteins are held in a partially autoinhibited state at the membrane *in vivo*, consistent with our prior in vitro observations (Stanishneva-Konovalova et al., 2016). The fact that loss of Nwk increases the frequency of patches while decreasing FM uptake suggests that the actin structures assembled in the *nwk* mutant are unproductive for synaptic vesicle endocytosis. These spurious patches could reflect non-specific actin assembly, perhaps due to unmasking of the Nwk ligand PI(4,5)P_2_ at the membrane and/or inappropriate activation of alternative WASp-dependent actin assembly pathways. Indeed, additional WASp activators such as Snx9 and Cip4/Toca-1 may play accessory roles in endocytic actin assembly (Almeida-Souza et al., 2018; Gallop et al., 2013), consistent with our finding that loss of presynaptic WASp led to a decrease in the total number of patches (**Fig 1E-G, Fig S1D-F**). Our data indicate that at the synapse, where endocytic machinery accumulates at high concentrations (Wilhelm et al., 2014) and recruitment appears uncoupled from activation, these layers of autoregulation are critical to constrain actin assembly generally.

Our findings on autoinhibition and clamping connect two prevailing models of the organization and function of the synaptic endocytosis machinery-preassembly and delivery. In the first model, preassembly of clathrin and accessory proteins is hypothesized to ensure fast endocytosis in response to synaptic vesicle fusion (Hua et al., 2011; Mueller et al., 2004; Wienisch and Klingauf, 2006). Here, Nwk autoinhibition provides a mechanism to assemble an inactive, yet poised endocytic apparatus. In the second model, endocytic machinery associates with the synaptic vesicle pool, providing a ready source or buffer of proteins that can be released to the plasma membrane upon calcium signaling or vesicle fusion (Bai et al., 2010; Denker et al., 2011; Gerth et al., 2017; Winther et al., 2015). Because Dap160/Intersectin can shuttle between the synaptic vesicle pool and the plasma membrane, is itself subject to autoregulation (Gerth et al., 2017), and can regulate other endocytic proteins (e.g. Dynamin, Nwk), it could serve as a single activator that couples the preassembly and delivery models.

### Coordinated relief of autoinhibition restrains and directs membrane-associated actin dynamics

Our *in vitro* data show that beyond functioning as a clamp, Nwk and Dap160 collaboratively activate WASp to promote robust actin assembly. Together with the defects we observed *in vivo* for actin dynamics and FM dye uptake, these data suggest that Dap160-Nwk-WASp interactions could serve as a coincidence detection mechanism to relieve autoinhibition of Nwk and promote productive actin assembly with other WASp activators. Coincidence detection mechanisms for F-actin assembly have been demonstrated in several models of endocytosis (Case et al., 2019; Sun et al.), suggesting that amplification of WASp membrane binding could drive robust actin patch assembly at synapses. Similarly, in human cells, the interaction between FCHSD2, Intersectin, and WASp promotes actin assembly and endocytic maturation (Almeida-Souza et al., 2018) or initiation (Xiao et al., 2018). The Dap160-Nwk module could act by directing and/or organizing actin assembly specifically at endocytic events, akin to the membrane directed actin assembly we observed *in vitro* (**Fig 6C**), and/or ensure that it is sufficiently robust for productive membrane remodeling (Akamatsu et al., 2020). Direct support for these models will require new analytical or imaging approaches to directly visualize the coupling of membranes and actin to the endocytic machinery, in order to distinguish spurious (due to unclamping) vs *bona fide* but underpowered endocytic actin assembly events.

### Physiological implications of autoregulatory mechanisms in the periactive zone

Our data suggest that the endocytic machinery can be deployed as clamped, primed, or activated complexes at different locations at the synapse. The next critical step will be to determine how physiological cues such as calcium influx or synaptic vesicle release might recruit or activate Dap160 or WASp to regulate transitions between these states. One intriguing possibility is that these mechanisms might enable an endocytic periactive zones to rapidly switch between different modes of endocytosis (e.g. ultrafast, conventional, or bulk) in response to a wide range of synaptic activity patterns (Gan and Watanabe, 2018). These endocytic regulatory mechanisms could also be locally poised to regulate, respond, or adapt to the specific release properties of nearby active zones (Akbergenova et al., 2018; Melom et al., 2013), and serve as novel points of control over synaptic plasticity and homeostasis.

## STAR Methods

### RESOURCE AVAILABILITY

#### Lead Contact

Further information and requests for resources and reagents should be directed to and will be fulfilled by the Lead Contact, Avital Rodal.

#### Materials Availability

All plasmids and fly lines generated in these studies are available upon request.

#### Data and Code Availability

All custom FIJI/ImageJ and Python scripts used to analyze the data in these studies are available upon request.

### EXPERIMENTAL MODEL DETAILS

#### Drosophila culture

Flies were cultured using standard media and techniques. All flies were raised at 25° C, with the exception of experiments using Dcr2; Dap160 RNAi or WASp RNAi, for which flies were raised at 29° C. See key resources table for all fly lines used and see Supplemental Table 1 for full genotypes for each experiment in this study.

### METHODS DETAILS

#### Cloning

UAS-Dap160 constructs were generated in pBI-UASC-mCherry (derived from (Wang et al., 2011), see (Deshpande et al., 2016)). Fragments were amplified from the genomic Dap160 locus with primers described in the key resources table. These transgenes were injected into flies (Rainbow Gene), using ΦC31-mediated integration at the Attp40 locus (Ni et al., 2008), to ensure that all constructs were in a similar genomic context. UAS-WASp-tev-myc was generated in pUAST (Brand and Perrimon, 1993) by inserting a TEV recognition site and 9 copies of the myc epitope tag at the 3’ end of the Wsp cDNA, and injected into w^1118^ flies at the Duke Model Systems Transgenic Facility (Duke University, Durham, NC).

#### Generation of AP2α::GFP^KI^

The vector pHD-sfGFP-dsRed was created using Gibson assembly by amplifying sfGFP from pScarlessHD-sfGFP-DsRed (gift from Kate O’Connor-Giles, Addgene plasmid # 80811) and inserting it in between the multiple cloning site and the first loxP site in the pHD-DsRed backbone (gift from Kate O’Connor-Giles, Addgene plasmid # 51434). 1 kb sequences upstream and downstream of the stop codon were amplified from the genomic locus of AP2α and inserted into pHD-sfGFP-dsRed using AarI and SapI, respectively, to create the HDR donor pMM007_pHD-AP2a-C-sfGFP-dsRed. The guide RNA GGAAATCTGCGATCTGTTGA was cloned into pU6-BbsI-chiRNA (gift from Melissa Harrison & Kate O’Connor-Giles & Jill Wildonger, Addgene plasmid # 45946 (Gratz et al., 2013); using BbsI to create pMM008_pU6-AP2a-chiRNA. 500 ng/uL HDR donor plasmid and 100 ng/uL gRNA plasmid were injected into vas-Cas9(III) flies (BDSC 51324, injections by BestGene). Correct integration of the transgene was validated by sequencing.

#### FM dye uptake

FM dye (FM1-43 in *dap160* experiments, or FM4-64FX in *nwk* experiments) uptake experiments were performed essentially as described (Ramachandran and Budnik, 2010; Verstreken et al., 2008). For fixed experiments (*nwk* mutants), larvae were dissected in groups of 4-6 (with each dish having at least two control larvae) in low-calcium HL3 (Stewart et al., 1994), and axons were cut to dissociate CNS input. For live imaging (*dap160* rescues), larvae were dissected, stained, and imaged in pairs, with one control (Dap160^FL^) and one experimental larva per dish. This temperature has been shown to exacerbate endocytic defects in some mutants, including *dap160* (Koh et al., 2004). Following extensive washing in Ca^++^ free saline, larvae were fixed in 4% paraformaldehyde in Ca^++^ free saline (for *nwk* experiments) or imaged live (for *dap160* transgene rescue experiments). Images of muscle 6/7 NMJs (abdominal segments 3-5) were acquired by confocal microscopy and FM dye intensity was measured within mCherry (in *dap160* experiments) or GFP (in *nwk* experiments)-labeled presynaptic masks, and intensities were normalized to dish-matched control larvae. For unloading experiments, larvae were analyzed individually. In all experiments, dye loading (4 μM) was performed in 90 mM KCl, 2 mM CaCl2 HL3 saline for five minutes at 36°C on a submerged metal block using prewarmed buffer. For unloading, larvae were stimulated for an additional 5’ with 90mM KCl 2mM CaCl2, washed extensively in Ca++ free HL3, then imaged and analyzed as for fixed larvae.

#### Immunohistochemistry

For analysis of NMJ morphology and protein localization, flies were cultured at low density at 25°C. Wandering third instar larvae were dissected in calcium-free HL3.1 saline (Feng et al., 2004) and fixed for 30 min in HL3.1 containing 4% formaldehyde. For analysis of NMJ overgrowth (satellite boutons), samples were stained with anti-HRP and anti-Dlg, and images were blinded before manual bouton counting. Boutons were counted on muscle 4 NMJs, abdominal segments 2-4, and satellite boutons were defined as any string of fewer than five boutons that branched from the main NMJ branch (O’Connor-Giles et al., 2008).

#### Western blots

*Drosophila* heads (10 pooled/genotype) were homogenized in 100 μL 2x Laemmli buffer. 10uL of extract per lane were fractionated by SDS/PAGE and immunoblotted with α-Dap160 (Roos 1998) and α-Tubulin antibodies (clone B-5-1-2, Sigma), and infrared-conjugated secondary antibodies (Rockland, Inc). Blots were analyzed on a Biorad Chemidoc system,

#### Imaging and analysis

Spinning-disk confocal imaging of *Drosophila* larvae was performed at room temperature (except *shi*^TS1^ experiments) on a Nikon Ni-E upright microscope equipped with 60× (NA 1.4) and 100× (NA 1.45) oil immersion objectives, a Yokogawa CSU-W1 spinning-disk head, and an Andor iXon 897U EMCCD camera. Images were collected using Nikon Elements AR software. For s*hi*^TS1^ GFP::actin imaging experiments, temperature was controlled using a CherryTemp temperature control unit (Cherry Biotech).

FRAP data (**Fig 3C**) were acquired on a Zeiss 880 with Airy Scan in super resolution acquisition mode, using a 63X NA1.4 objective. Single Z-slices through the middle of individual boutons were acquired at 4Hz for 90sec, with manual focus adjustment. Following acquisition of two or three initial Z-stacks to assess prebleach intensity, <~20% of individual boutons were photobleached by the 488 laser at .6 intensity and scan speed of 6. Intensities of background, unbleached, and bleached ROIs were acquired manually using FIJI, and bleached area was normalized to prebleach and unbleached ROIs (to correct for imaging-induced photobleaching), and analyzed with GraphPad Prism.

Confocal imaging of GUVs and cell-sized water droplets was conducted at room temperature on a Marianas spinning disk confocal system (3I, Inc, Denver,CO), consisting of a Zeiss Observer Z1 microscope equipped with a Yokagawa CSU-X1 spinning disk confocal head, a QuantEM 512SC EMCCD camera, PLAN APOCHROMAT 63X or 100x oil immersion objectives (n.a. 1.4), a Photonics Instruments Micropoint photo-ablation device, and Slidebook software.

3D-Structured Illumination Microscopy (SIM) was performed on a Nikon N-SIM E system (consisting of an inverted Eclipse Ti-E microscope, 100x (n.a 1.45) oil-immersion objective, and a Hamamatsu OrcaFLASH4 sCMOS camera). Channel alignment was calculated for each imaging session using tetraspeck beads (Invitrogen cat no. T-7284). Images were collected at room temperature with a regime of 3 grid orientations and 5 phases and were reconstructed using Nikon Elements software, using a theoretical, ideal OTF generated by the software. Super-resolution images of protein localization in live samples were acquired with a Zeiss 880FAS microscope in fast Airyscan mode with a 63X (NA1.4) oil immersion objective, using Zen Blue software.

#### Analysis of actin dynamics at the NMJ

Spinning disk confocal time series were acquired at 15 stacks/min, except for experiments in Fig 6D-F, which were acquired at 60 stacks/min. A maximum intensity projection was made of each time point, movies were registered using the FIJI plugin StackReg, and analyzed by Patchtracker (Berro and Pollard, 2014) as follows: The threshold for patch detection was normalized to the mean probe intensity in the presynaptic area (threshold=Probe Mean * 0.40). All other settings for patch detection and tracking were default: Estimated patch diameter .6μm, median filter=false, subpixel detection=true. Linking max distance=.5μm, Gap-closing distance=.5 μm, Gap-closing frame gap=0. For .25Hz imaging experiments, patches between 16-356s could be detected. For 1Hz imaging experiments, patches between 4-139s could be detected. Because this analysis rejects a significant number of detected patches due to tracking defects or tracking path overlap, we estimated the true patch frequency as follows: We combined detections from 0.25Hz and 1Hz imaging experiments by averaging the frequencies over the shared detection range (20-150 seconds) and adding the lower and higher duration patches that were specific to each imaging regime (4-16 seconds for 1Hz and 150-360 seconds for 0.25Hz). Then we ‘corrected’ for rejected tracks, and considered the lower bound of the estimate to be the actual, uncorrected merged frequency of detection (2.8 patches/10μm/min) and the upper bound to be include every rejected track (10.3 patches/10 μm/min).

Actin dynamics were also analyzed by measuring intensity variation over time over the entire NMJ, i.e. without thresholding or particle tracking. We measured this by extracting the intensity value for each pixel over time and calculating the coefficient of variation (CoV, StdDev/Mean) for each pixel. We estimated the percentage of ‘highly variant’ pixels by thresholding these values using Li (Li and Tam, 1998) and Moments (Tsai, 1995) algorithms. While these two algorithms gave different estimates of the fraction of NMJs covered by highly variant pixels, both indicated the same relationship between genotypes. To validate this approach we created synthetic data using a custom FIJI script, with a spatial and temporal scale that matched our *in vivo* imaging, and in which we varied parameters expected to impact this metric (signal intensity, noise level, fraction of dynamic pixels, dynamics frequency, dynamics duration, dynamics amplitude), and subjected the synthetic data to the same CoV over time analysis.

#### Intensity and Colocalization analysis

For intensity and colocalization, the presynaptic region was masked in 3D using a presynaptically enriched label: either HRP (Fig 3E, S4B), Nwk (Fig 1C, S1A, 3F), Dap160 (S1B), or Lifeact::ruby (Fig 2B). For mask generation, images were subjected to a gaussian blur filter and thresholded by intensity. Blur radius and the specific threshold algorithms used were empirically optimized for each experiment to consistently and accurately reflect the presynaptic area in control and mutant groups (and the same settings were used for all NMJs within any given experiment). Signal intensities were measured in 3D using a FIJI script, and colocalization analysis was performed in 3D on Airyscan or SIM reconstructed image stacks using the Coloc2 plugin for ImageJ (https://imagej.net/Coloc_2). For all images, background was subtracted using the rolling ball method with a radius of 50 pixels.

### In vitro assays

#### Protein purification

Dap160^SH3C^ and Dap160^SH3CD^ were amplified from Dap160 Isoform A and cloned into pTrcHisA (see STAR methods for details). N-terminally His-Xpress–tagged proteins (Nwk^1-633^, Nwk^1-731^, Nwk^607-731^, Nwk^1-428^, Wsp^143-529^, Dap160^SH3C^, Dap160^SH3CD^), were purified as described previously (Becalska et al., 2013; Kelley et al., 2015; Rodal et al., 2008; Stanishneva-Konovalova et al., 2016). In brief, proteins were purified from BL21(DE3) *Escherichia coli* using cobalt or nickel columns, followed by ion exchange and gel filtration into 20 mM Tris pH 7.5, 50 mM KCl, 0.1 mM EDTA, and 0.5 mM DTT. GST fusions (Dap160^SH3CD^, Dap160^SH3C^, Dap160^SH3D^) were amplified from Dap160 isoform A and cloned into pGEX4t (see STAR methods for details). Proteins were purified with glutathione agarose (Thermo Scientific, Waltham MA) in 20 mM Tris 7.5, 20 mM KCl, and 0.5 mM dithiothreitol (DTT) supplemented with protease inhibitors (P2714 (Sigma-Aldrich, St. Louis, MO) and 0.5 mg/mL pepstatin A). Arp2/3 complex was purchased from Cytoskeleton, Inc. Actin was purified from from acetone powder (Spudich and Watt, 1971) generated from frozen ground hind leg muscle tissue of young rabbits (PelFreez, Rogers, AR).

#### Coprecipitation assays

Coprecipitation with GST-tagged proteins were conducted as described previously (Kelley et al., 2015): Concentrations of GST fusions on beads were normalized using empty beads and bead volume was restricted to two-thirds of the total reaction volume. GST fusions were incubated with agitation with His-tagged target proteins at room temperature for one hour in binding buffer (20 mM Tris pH 8.0, 20 mM KCl, 0.5 mM DTT). For salt sensitivity experiments, the indicated concentrations of NaCl were used in place of KCl in the binding buffer. Beads were then pelleted and washed once with buffer after removing the supernatant. Pellets and supernatants were then boiled in Laemmli Sample Buffer and fractionated by SDS-PAGE, followed by Coomassie staining or immunoblotting as noted in figure legends, followed by imaging and analysis on a LICOR Odyssey device.

#### Liposome cosedimentation

Lipid cosedimentation assays were conducted as described previously (Becalska et al., 2013). In brief, liposomes were swelled from dried lipid films in 20 mM Hepes pH 7.5 and 100 mM NaCl. Specific lipid compositions are indicated in the figure legends. Proteins were then mixed with 1 mg/mL liposomes, incubated for 30 min at room temperature, and then pelleted for 20 min at 18,000 × g at 4 °C. Pellets and supernatants were then denatured in Laemmli sample buffer and fractionated by SDS/PAGE, followed by Coomassie staining, and imaging and analysis on a LICOR Odyssey device.

#### GUV decoration

GUVs were generated by gentle hydration. Briefly, 10 μL of 10 mg/mL lipids dissolved in 19:1 chloroform:methanol were dried under vacuum, then swelled in 300 μL of 5 mM HEPES 300 mM sucrose, pH 7.5 overnight at 70°C. GUVs were imaged on a Marianas spinning disk confocal system (see above). 3 μL GUVs were diluted into 5 mM HEPES 150 mM KCl pH7.5, incubated with protein as noted in figures, and imaged using a 100X/ N.A. 1.4 objective in multiwell slides (Lab-Tek) precoated with 1 mg/mL BSA. After a 30 min incubation, 1% agarose in 5 mM HEPES 150 mM KCl pH 7.5 was added (final agarose concentration .5%) to limit GUV mobility. Images were analyzed by line tracing intensity profiles across a medial optical section of GUVs.

#### Actin assembly in droplets

Lipids (97.5% DPHPC [1,2-diphytanoyl-sn-glycero-3-phosphocholine], Avanti Polar Lipids) and 2.5% DPHPC:PI(4,5)P_2_) were mixed in chloroform, dried under vacuum, and rehydrated to 23 mM (20 mg/ml) in decane. The indicated proteins were added to the lipid mix at a 1:50 volume ratio and pipetted vigorously until cloudy before imaging by spinning disk confocal microscopy.

#### Pyrene-actin assembly

Rabbit muscle actin [5% (mol/mol) pyrene-labeled] was gel-filtered, prespun at 90,000 x g, exchanged from Ca^2+^ to Mg^2+^, and assembled at a final concentration of 2.5 μM as described previously (Moseley et al., 2006). Proteins were preincubated with 74 μg/mL liposomes or control buffer for 30 min before actin assembly reactions. Assembly was monitored with a spectrofluorometer (Photon Technology International) using an excitation wavelength of 365 nm and an emission wavelength of 407 nm. Rates were calculated from slopes of curves in the linear range, and curves were plotted using GraphPad Prism software.

### QUANTIFICATION AND STATISTICAL ANALYSIS

Graphs were prepared and statistical analyses performed using Graphpad Prism software. For normally distributed data, comparisons were made using either T-test or ANOVA with posthoc Bonferroni’s multiple comparisons test. For non-normally distributed data, comparisons were made using either Mann-Whitney U test, or Kruskal-Wallis test with posthoc Dunn’s test. Comparison of patch duration distributions was performed using a Kolmogorov-Smirnoff test. Please see Supplemental Table 1 for each statistical test performed for each experiment presented in this study. All data are shown as the mean +/− SEM.

## SUPPLEMENTAL MEDIA

**Supplemental Movie 1.** Dynamics of actin patches labeled by complementary reporters.

**Supplemental Movie 2.** Loss of *nwk* increases the frequency of brief actin patches.

**Supplemental Movie 3.** AP2α and Lifeact::Ruby partially colocalize.

## Author Contributions and Notes

Conceptualization S.J.D, C.F.K, E.M.M, A.A.R. Methodology S.J.D, C.F.K, E.M.M, T.F., A.A.R. Software S.J.D. Formal Analysis S.J.D, C.F.K, M.F.M, E.M.M. Investigation S.J.D, C.F.K, E.M.M, T.L, M.F.M, B.E. Resources C.F.K, E.M.M, T.L, M.M, M.K. Data curation S.J.D, C.F.K, E.M.M, A.A.R. Writing – Original Draft S.J.D, C.F.K, A.A.R. Writing – Review & Editing S.J.D, C.F.K, B.E, M.M, M.K, A.A.R. Visualization S.J.D, C.F.K, A.A.R. Supervision M.K, A.A.R. Project Administration A.A.R. Funding Acquisition M.K, A.A.R.

The authors declare no conflict of interest.

This article contains supporting information online.

## Acknowledgments

The authors would like to thank Bruce Goode for actin reagents and advice, Julien Berro for particle tracking advice, Troy Littleton and Oleg Shupliakov for helpful discussions, Graeme Davis for anti-Dap160 antibody, and the Bloomington *Drosophila* Stock Center (Indiana University, Bloomington, IN, NIH P40OD018537) for providing fly stocks. This work was supported by a Basil O’Connor Scholar Award from the March of Dimes and a Pew Scholar award (AAR); by R01 NS116375 (AAR and TGF); by the Brandeis NSF MRSEC, Bioinspired Soft Materials (NSF-DMR 1420382); by T32 NS007292 (SJD), T32 GM007122 (CFK), and R90 DA03243501 (MFM); and by the Swiss National Science Foundation (grant 310030B_182825) and NCCR Chemical Biology funded by the SNSF (MK, MM).

