## Supplemental Material for "An autoinhibitory clamp of actin assembly constrains and directs synaptic endocytosis"

**Supplemental Movie 1. Dynamics of actin patches labeled by complementary reporters**. Actin reporters driven by C155-Gal4 (single timepoints shown in **Fig. S2C**). Timelapse Z-stacks were acquired by spinning disc confocal microscopy at 4 sec/frame. Movie shows MaxIPs played at 15X live speed. The predominant structures are transient patches (quantified in **Fig. 1B-D**), though we also observed much less frequent structures that exhibited significant lateral mobility or resembled 'comet tails', and other more stable, cable-like structures. Scale bar is 5 µm. Associated with **Figs 1, S1**.

**Supplemental Movie 2 Loss of nwk increases the frequency of brief actin patches.** MaxIPs of spinning disc confocal timelapses of control (left) and $nwk^{1/2}$ (right) muscle 6/7 NMJs acquired at 1Hz (playback 18 fps). Nwk mutants exhibit spurious synaptic F-actin assembly. Scale bar is 5µm. Associated with **Fig 6D**.

**Supplemental Movie 3. AP2α and Lifeact::Ruby partially colocalize**. Timelapse Z-stacks were acquired by Airyscan imaging at 4 sec/frame.of endogenously tagged AP2α::GFP and actin patches labeled by Lifeact::Ruby driven by C155-Gal4 (single timepoints shown in **Fig. 2I**). Movie shows MaxIP played at 16X live speed. Associated with **Fig 7**.

Figure 1 S1

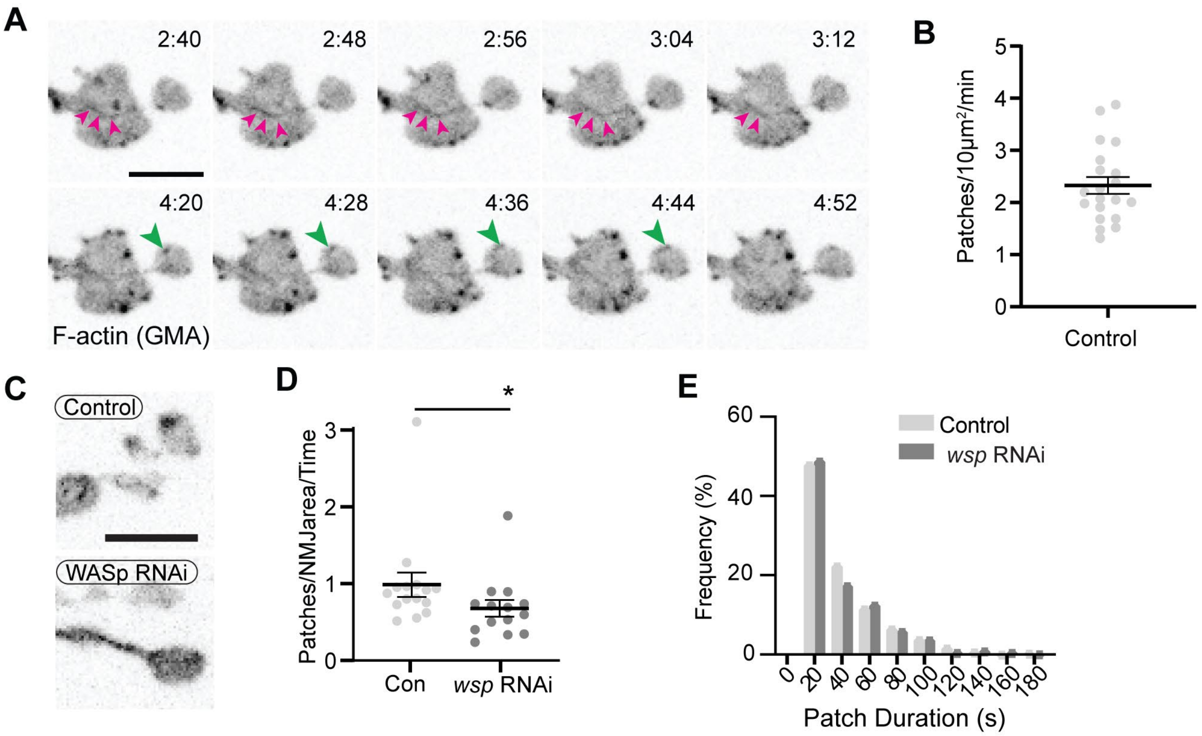

**Figure 1 Supplement 1. Additional characterization of actin patches.** (A) Image series from **Movie 1** highlighting actin cables (magenta arrowheads) and in addition to dynamic patches (large green arrowheads). (B) Quantification of patch frequency in NMJs expressing GMA and imaged at 1Hz. Data are from same experiment as controls in **Fig 6A-C** (C) Representative MaxIPs of single spinning disk confocal microscopy time points of neuronally expressed GMA in control and WASp RNAi (expressed in neurons via C155-GAL4) NMJs. (D,E) Quantitative analysis of patch frequency and patch duration distribution (imaged at 0.25 Hz) shows that presynaptic WASp RNAi recapitulates the decreased patch frequency observed at *WASp* mutant NMJs. Histogram bins are 20 sec; X axis values represent bin centers. Scale bars in A and C are 5 µm. Graphs show mean +/- s.e.m.; n represents NMJs. Associated with **Fig 1, Movie 1**.

Figure 3 S1

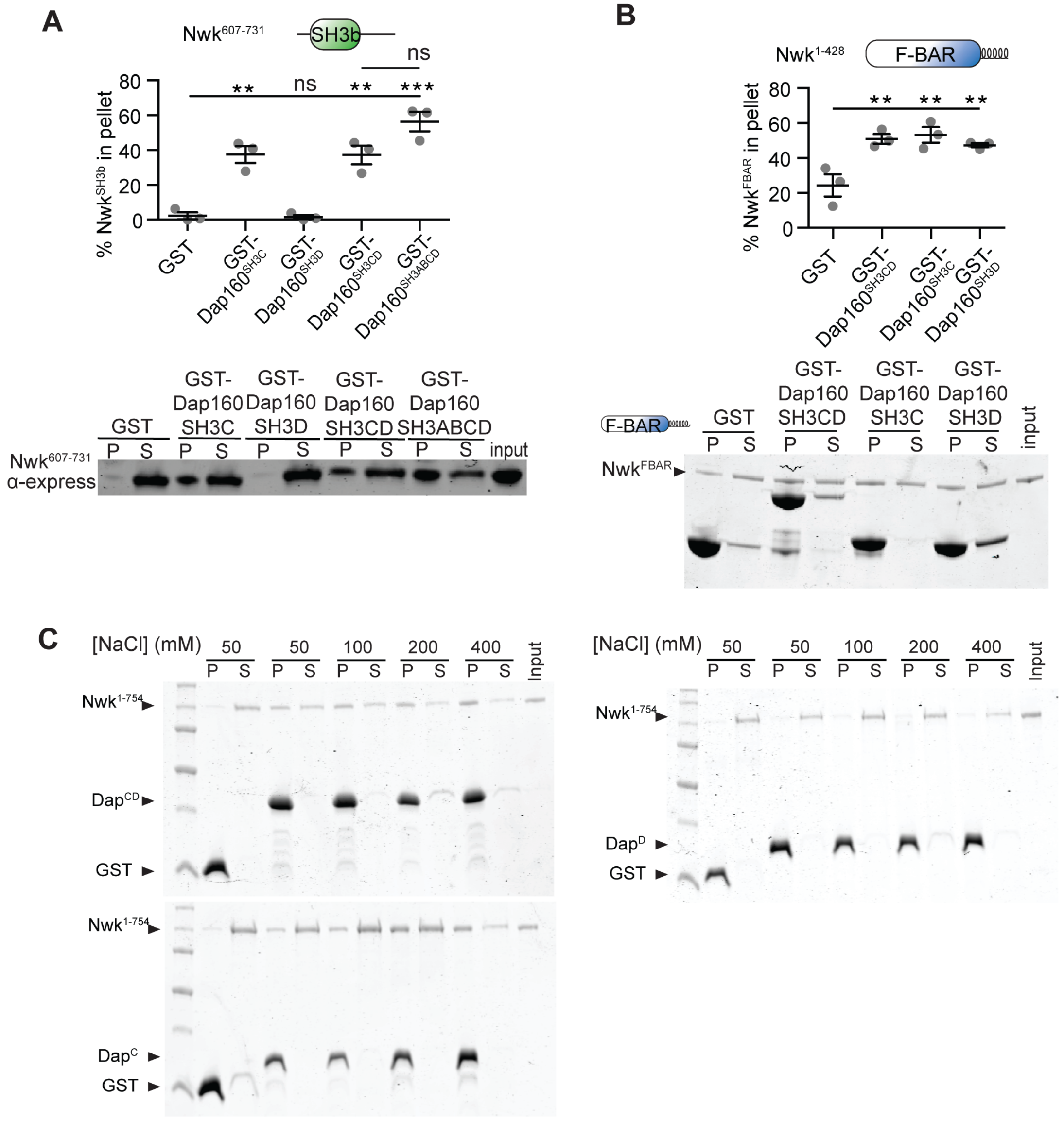

**Figure 3 Supplement 1.** (A-B) GST fusion proteins were immobilized on glutathione agarose and incubated with the indicated purified proteins. Pellets and supernatants were fractionated by SDS-PAGE, immunoblotted (A) or Coomassie stained (B) and quantified by densitometry. (A) $Nwk^{SH3b}$ interacts directly with $Dap160^{SH3C}$, $Dap160^{SH3CD}$ or $Dap160^{SH3ABCD}$, but not with $Dap160^{SH3D}$ alone. [$Nwk^{SH3b}$]=7μM. (B) The Nwk F-BAR domain interacts directly with $Dap160^{SH3}$, $Dap160^{SH3D}$, and $Dap160^{SH3CD}$. [$Nwk^{F\text{-}BAR}$]=1.5μM, [GST-$Dap160^{SH3CD}$]=8.5μM, [GST-$Dap160^{SH3C/D}$]=11μM. (C) Representative Coomassie-stained gels for **Fig 3B.**

Figure 3 S2

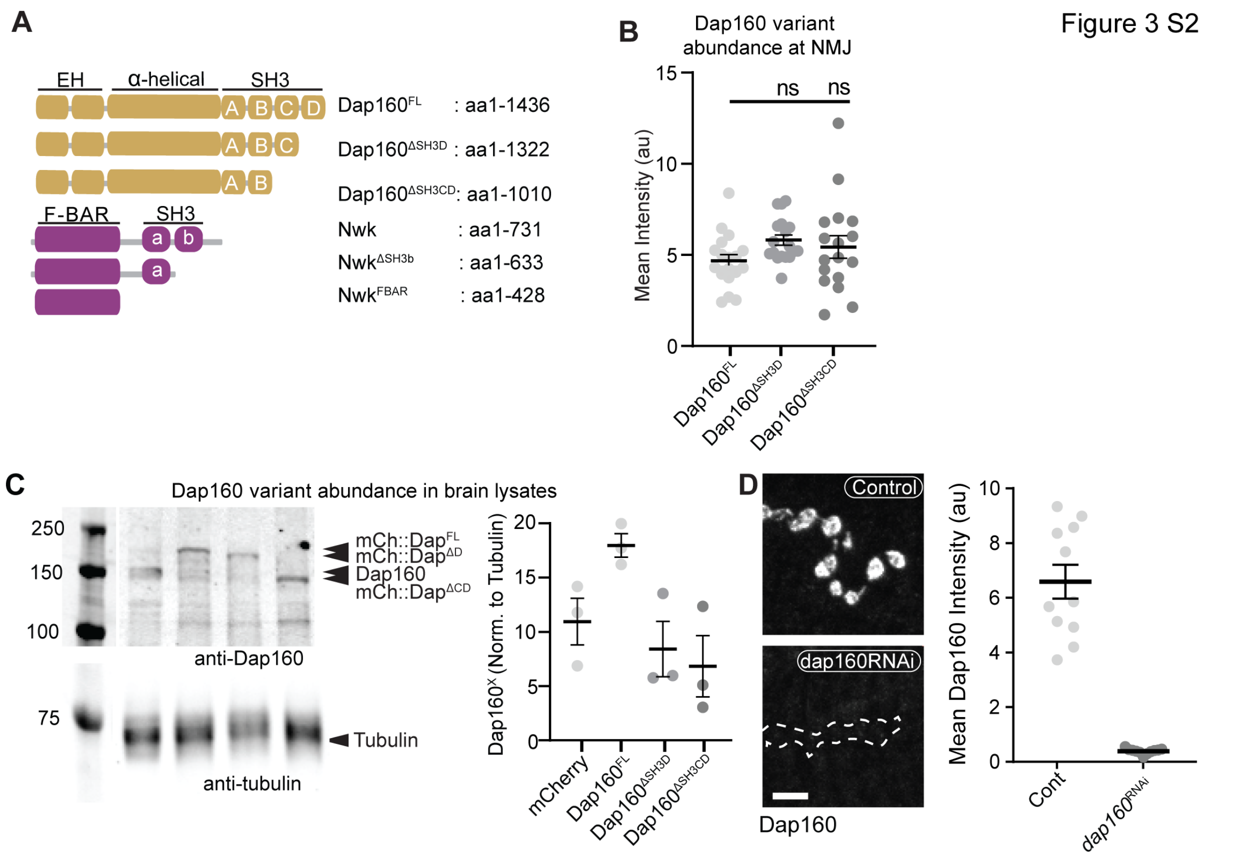

**Figure 3 Supplement 2.** (A) Schematic of Dap160 rescue transgenes used *in vivo* and Nwk fragments used *in vitro* experiments. (B) Quantification of Dap160 variant transgene expression (mCherry signal) at NMJs from dataset in **Fig 3E**. All transgenes were expressed in neurons by C155-GAL4 in a *dap160* null background (*dap160*$^{\Delta 1/Df}$). (C) Representative Western blots (left) and quantification (right) of Dap160 rescue transgene expression in *Drosophila* adult head extracts. (D) MaxIPs (left) and quantification (right) of spinning disc confocal micrographs of control and *dap160* RNAi expressing NMJs showing knockdown of Dap160 protein levels, using the same conditions as in **Fig 3D**. Associated with **Fig 3**.

Figure 5 S1

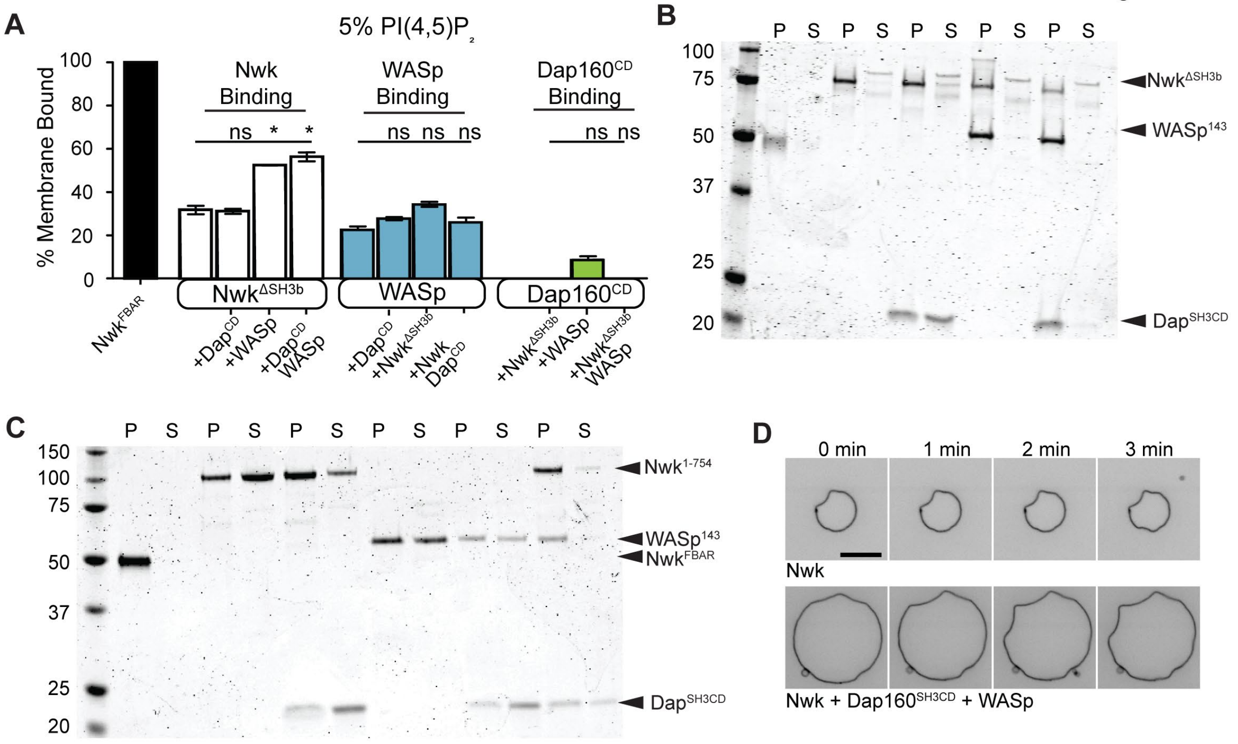

**Figure 5 Supplement 1.** (A) Cosedimentation assay between the indicated purified proteins and liposomes composed of [mol% = DOPC/DOPE/DOPS/PI(4,5)$P_2$ = 75/15/5/5]. Dap160$^{SH3CD}$ is unable to enhance membrane binding of Nwk$^{\Delta SH3b}$ (which lacks the autoinhibitory and Dap160-binding SH3b domain), or WASp. Conversely, Nwk$^{\Delta SH3b}$ is unable to promote membrane recruitment of Dap160$^{SH3CD}$. Quantification from Coomassie-stained gels represents the mean fraction of total protein that cosedimented with the liposome pellet, ± SEM. [Nwk$^{1\text{-}xxx}$] = 2µM, [Dap160] = 6µM. Graph shows mean +/- s.e.m. (B) Representative Coomassie-stained gel from (A). (C) Representative Coomassie-stained gel from **Fig 5C**. (D) Confocal timelapse of deformation of GUVs (10% PI(4,5)$P_2$, labeled with <1% TopFluor-PE) decorated with the indicated proteins [Nwk$^{1\text{-}xxx}$] = 250 nM, [WASp]=250 nM, [Dap160] = 1.2 5µM. The Nwk-Dap160-WASp complex retains the same membrane remodeling activity as Nwk alone. Associated with **Fig 5**.

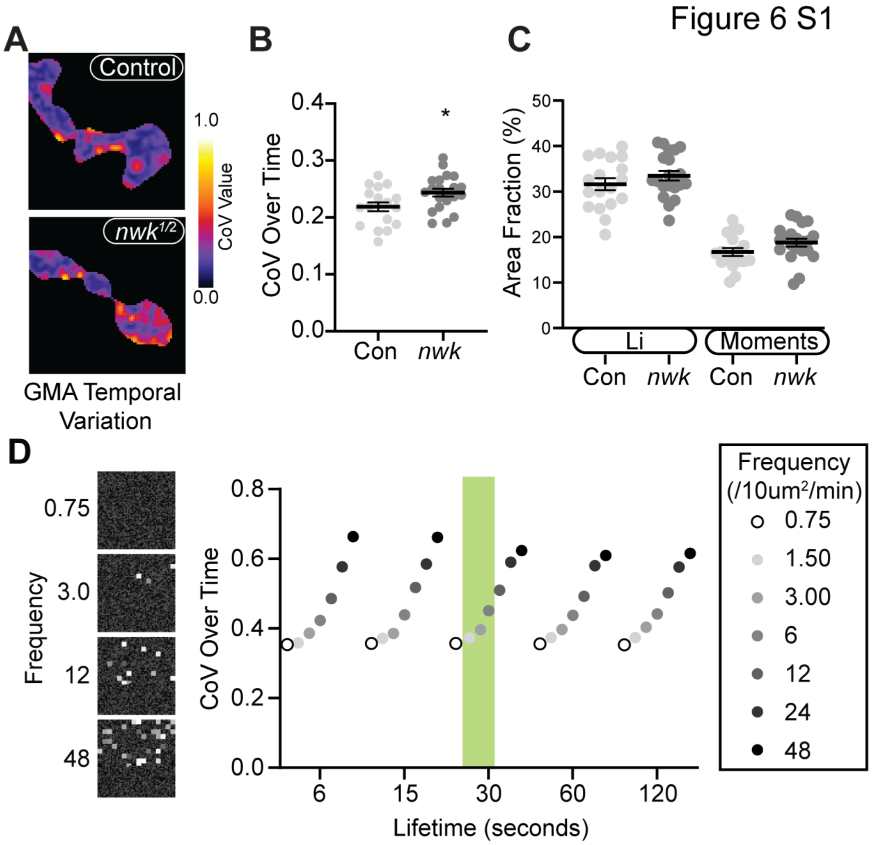

**Figure 6 Supplement 1.** (A) CoV projections of time-lapse movies of GMA dynamics in control and nwk mutant NMJs (analysis of same dataset as shown in 6A-C; different representative images shown). (B) Average temporal CoV values across NMJs. *nwk* mutants exhibit greater temporal variation in GMA signal than controls. (C) Quantification of the fraction of NMJ area covered by 'high' CoV pixels (identified by automated thresholding using either Li (Li and Tam, 1998) or Moments (Tsai, 1995) algorithms). *nwk* mutants show no difference in the fraction of high CoV pixels, suggesting that actin dynamics are confined to a restricted region of the synapse, and vary more over time rather than space. (D) Example of results from temporal CoV analysis of synthetic data, in which the frequency and lifetime were systematically varied. (Left) Representative images of synthetic data created using a custom FIJI script. (Right) Quantification of CoV over time. Shaded green region indicates values that most closely match in vivo measurements drawn from Patchtracker particle-based analysis. In this regime, our model indicates that the *nwk* mutant effect size in panel B corresponds to a 43% increase in patch frequency, slightly higher than the particle-based measurement of 28%.

Figure 7 S1

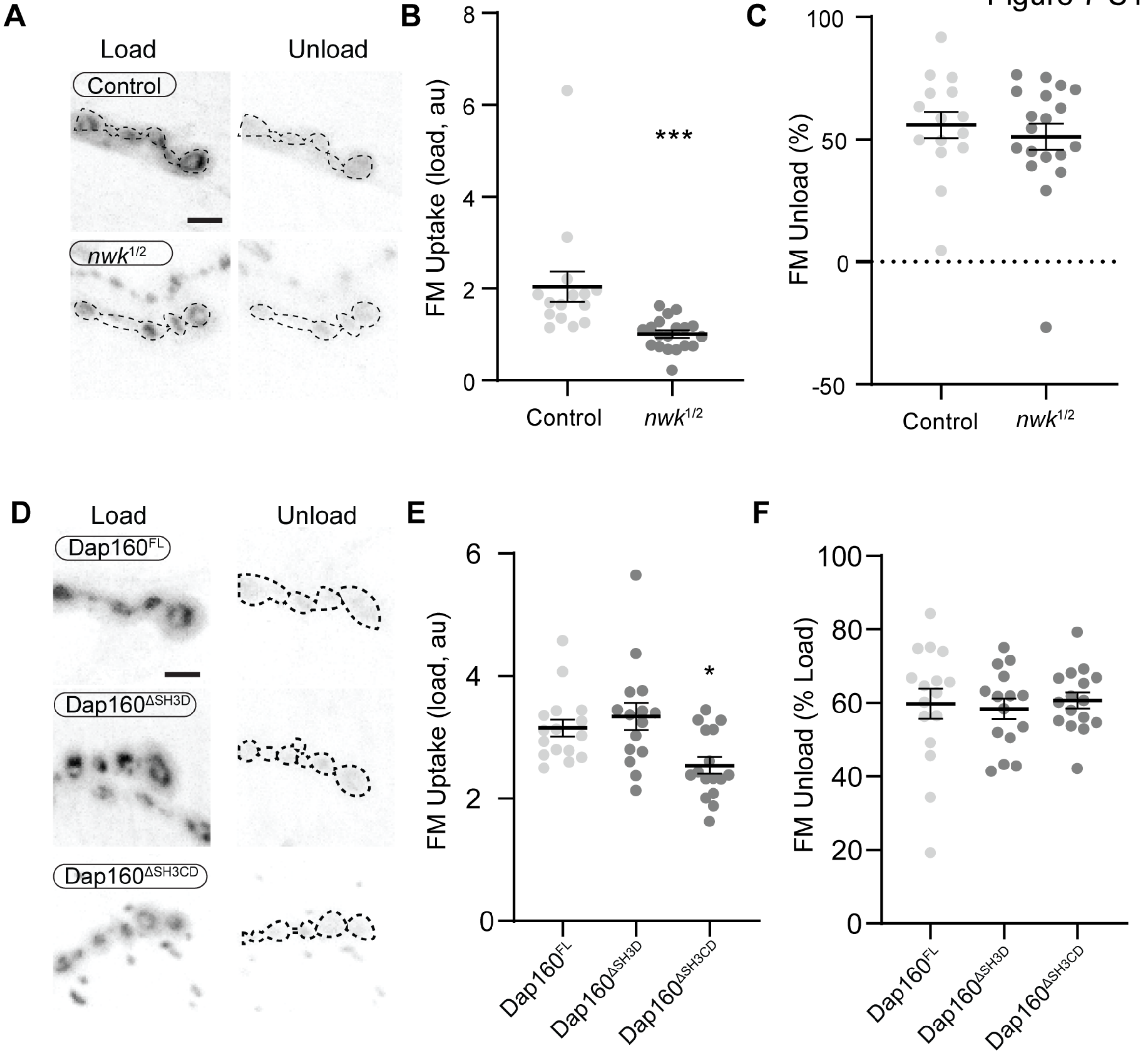

**Figure 7 Supplement 1.** Nwk and Dap160SH3 mutants do not disrupt FM dye unloading. Loading and unloading of FM dyes in *nwk* and *dap160*$^{SH3}$ mutants by stimulation in 90mM KCl + 2mM $CaCl_2$. (A,D) MaxIPs of spinning disc confocal stacks acquired after loading (left) and unloading (right) of FM dyes. (B-C) Quantification of FM4-64 loading (B) and unloading (C) in *nwk* mutants. (E-F) Quantification of FM1-43 loading (E) and unloading (F) in *dap160* domain mutants.

Figure 7 S2

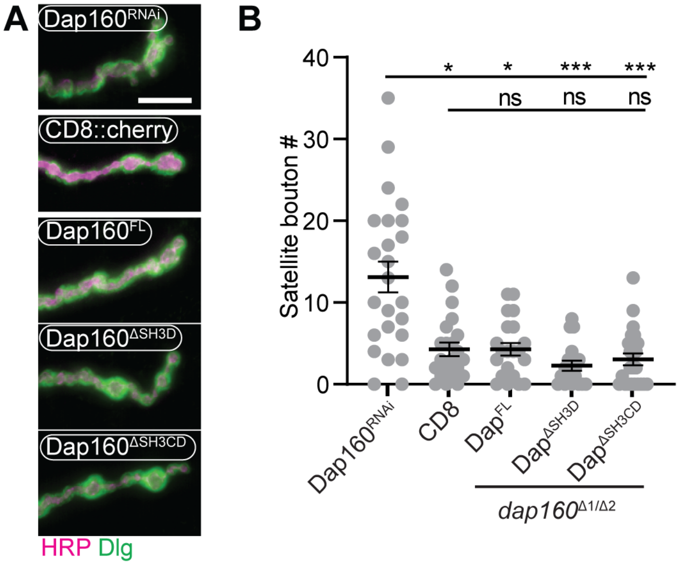

**Figure 7 Supplement 2.** (A-B) All Dap160 transgenes rescue *dap160* satellite bouton phenotype. (A) MaxIPs of epifluorescence micrographs of muscle 4 NMJs stained for anti-HRP (magenta) and anti-Dlg (green). $Dap160^{RNAi}$ expressing NMJs exhibit many satellite boutons, which are restored to normal levels by presynaptic expression of $Dap160^{FL}$, $Dap160^{\Delta SH3D}$, or $Dap160^{\Delta SH3CD}$ (in $dap160^{\Delta 1/Df}$ larvae). (B) Quantification of satellite boutons. Graph shows mean +/- s.em.; n represents NMJs. Scale bar in (A) is 10 μm. Associated with **Fig 7**.

**Supplemental Table 1. Summary of genotypes and statistics for all experiments in this study.**

| Figure/<br>Experiment | Genotype/Conditions | N | Statistical Test(s) |
|---|---|---|---|
| 1A-D<br>GFP::Actin<br>GMA<br>Lifeact::Ruby | <br>C155-GAL4/+ or Y; UAS-GFP::actin/+<br>C155-GAL4/+ or Y; UAS-GMA/+<br>C155-GAL4/+ or Y; UAS-lifeact::Ruby/+ | <br>823 patches/9 NMJ/5 larvae<br>819 patches/15 NMJ/6 larvae<br>363 patches/7 NMJ/3 larvae | NA |
| 1E-F<br>Arp3 | <br>C155-GAL4/UAS-Arp3::GFP; UAS-lifeact::Ruby/+ | <br>13 NMJ/3 larvae | NA |
| 1G-I<br>Control<br>WASp | <br>C155-GAL4/+ or Y; UAS-GMA/+<br>C155-GAL4/+ or Y; wsp1,e,UAS-GMA/wsp1,e | <br>832 patches/18 NMJ/8 larvae<br>532 patches/15 NMJ/6 larvae | Kolmogorov-Smirnoff (G) Welch's t test (H) |
| S1A-B | C155-GAL4/+ or Y; UAS-GMA/+ | 1606 patches/20 NMJ/8 larvae | Kolmogorov-Smirnoff |
| S1C-E<br>Control<br>WASp | <br>C155-GAL4/Y; UAS-GMA/UAS-luciferaseRNAi<br>C155-GAL4/Y; UAS-GMA/UAS-WASpRNAi | <br>709 patches/15 NMJ/6 larvae<br>286 patches/14 NMJ/5 larvae | Kolmogorov-Smirnoff (H)<br>Mann-Whitney (I)<br>t test (J) |
| 2A | Nwk$^{MIMICGFP}$ | | |
| 2B | Vglut-GAL4/Y; CD8::RFP/+; Nwk::GFP/+ | | |
| 2C-D | C155-GAL4/Y; UAS-WASp::myc/+; UAS-GMA/+ | 14 NMJ/3 larvae | NA |
| 2E-F | C155-GAL4/Y; UAS-WASp::myc/+; UAS-GMA/+ | 12 NMJ/3 larvae | NA |
| 2G-I<br>AP2 | <br>C155-GAL4/+; UAS-lifeact::Ruby/AP2::GFP | <br>14 NMJ/3 larvae | NA |
| 3B<br>GST-X<br>Nwk$^{1-731}$ | <br>5µg<br>3µg | 3 tech replicates per lane | NA |
| 3B<br>GST-CD<br>Nwk$^{1-428}$<br>Nwk$^{1-731}$ | <br>1.6µM CD, 1.2µM C/D<br>1.5µM<br>0.8µM | 3 tech replicates per lane | NA |
| 3D<br>Dap$^{FL}$<br>Dap$^{\Delta D}$<br>Dap$^{\Delta CD}$<br>Dap$^{RNAi}$ | rescues=C155-GAL4/+; dap160$^{\Delta 1}$/Df3450; X<br>X=UAS-Dap$^{FL}$::mCherry/+<br>X=UAS-Dap$^{\Delta D}$/+<br>X=UAS-Dap$^{\Delta CD}$/+<br>C155-GAL4, UAS-Dcr2/Y; UAS Dap160-RNAi/+ | <br>18 NMJ/3 larvae<br>18 NMJ/3 larvae<br>17 NMJ/3 larvae<br>16 NMJ/3 larvae | ANOVA+Tukey's multiple comparison test |
| 3E<br>Dap$^{FL}$<br>Dap$^{\Delta D}$<br>Dap$^{\Delta CD}$ | rescues= C155-GAL4/+; dap160$^{\Delta 1}$/Df3450; X<br>X=UAS-Dap$^{FL}$::mCherry/+<br>X=UAS-Dap$^{\Delta D}$/+<br>X=UAS-Dap$^{\Delta CD}$/+ | <br>17 NMJ/3 larvae<br>18 NMJ/3 larvae<br>17 NMJ/3 larvae | Kruskal-Wallis + Dunn's multiple comparison test |
| 3 S1A<br>Nwk$^{607-731}$ | <br>7µM | | |
| 3 S1B<br>GST-X<br>Nwk$^{1-428}$ | <br>.375mg/mL (CD=8.5µM, C/D=11µM)<br>1.5µM | 3 tech replicates per lane | ANOVA+Tukey's multiple comparison test |

| **Figure/ Experiment** | **Genotype/Conditions** | **N** | **Statistical Test(s)** |
|---|---|---|---|
| 3S1C | Same experiment as Figure 3B | Same experiment as Fig 3B | ANOVA+Tukey's multiple comparison test |
| 3S2B | Same experiment as Figure 3D | Same experiment as Fig 3D | ANOVA+Tukey's multiple comparison test |
| 3 S2C<br>mCh<br>$Dap^{FL}$<br>$Dap^{\Delta D}$<br>$Dap^{\Delta CD}$ | <br>C155-GAL4/+; UAS-CD8::RFP/+<br>X=UAS-$Dap^{FL}$::mCherry/+<br>X=UAS-$Dap^{\Delta D}$/+<br>X=UAS-$Dap^{\Delta CD}$/+ | <br>10 pooled brains/replicate,<br>3 biological replicates/group | NA |
| 3 S2D<br>Control<br>Dap160RNAi | <br>vglut/Y; Nwk::$GFP^{MiMIC}$/UAS-luciferaseRNAi<br>vglut/Y; UAS-dcr2/+; Nwk::$GFP^{MiMIC}$/$Dap160^{RNAi}$ | <br>11NMJ/3larvae<br>11NMJ/3larvae | t test |
| 4A<br>Actin<br>Arp2/3<br>WASp<br>Nwk1-731<br>DapCD<br>DapC | <br>2μM<br>50nM<br>50nM<br>500nM<br>2μM<br>2μM | <br>(1) 2 replicates<br>(2) 2 replicates<br>(3) 3 replicates<br>(4) 3 replicates<br>(5) 5 replicates<br>(6) 2 replicates | ANOVA+<br>Tukey's multiple comparison test |
| 4B<br>Actin<br>Arp2/3<br>WASp<br>Nwk1-731<br>DapCD<br>PI(4,5)$P_2$ | <br>2μM<br>50nM<br>50nM<br>100nM<br>500nM<br>2μM 10% PI(4,5)P2 liposomes | <br>(1) 2 replicates<br>(2) 2 replicates<br>(3) 3 replicates<br>(4) 3 replicates<br>(5) 3 replicates | ANOVA+<br>Tukey's multiple comparison test |
| 4C<br>$Nwk^{1-633}$<br>$Nwk^{1-731+}Dap^{CD}$ | 2μM OG-actin, 50nM Arp2/3, 50nM WASp<br>500nM $Nwk^{1-633}$::SNAP549<br>500nM $Nwk^{1-731}$::SNAP549+2μM $Dap160^{SH3CD}$ | <br>41 droplets<br>22 droplets | NA |
| 5A<br>$Nwk^{1-428}$<br>$Nwk^{1-754}$<br>$Dap^{CD}$ | DOPC/DOPE/DOPS/PI(4,5)$P_2$ = 70/15/5/10<br>3μM<br>1.125μM<br>1.69-6.75μM | 3 tech replicates per group | ANOVA+<br>Tukey's multiple comparison test |
| 5B<br>$Nwk^{1-XXX}$<br>$Dap160^{X}$ | DOPC/DOPE/DOPS/PI(4,5)$P_2$ = 80-x/15/5/x<br>2μM<br>6μM | 3 tech replicates per group, except 2 replicates for:<br>2.5%PIP2-$Nwk^{1-428}$<br>2.5%-$Nwk^{1-731}$+$Dap^{SH3C}$ | ANOVA+<br>Tukey's multiple comparison test |
| 5C<br>$Nwk^{1-731}$<br>WASp<br>$Dap160^{CD}$ | DOPC/DOPE/DOPS/PI(4,5)$P_2$ = 70/15/5/10<br>1μM<br>1μM<br>3μM | 3 tech replicates per group | ANOVA+<br>Tukey's multiple comparison test |
| 5D<br>$Nwk^{1-731}$<br>Nwk/WASp/Dap | 5% PI(4,5)$P_2$ GUVs<br>500nM $Nwk^{1-731}$::SNAP549<br>250nM $Nwk^{1-731}$::SNAP549, 250nM WASp, 1.25μM $Dap160^{SH3CD}$ | Representative from:<br>11 GUVs imaged<br>12 GUVs imaged | NA |
| 5E-F<br>Control<br>Dap160RNAi | <br>vglut/Y; Nwk::$GFP^{MiMIC}$/UAS-luciferaseRNAi<br>vglut/Y; UAS-dcr2/+; Nwk::$GFP^{MiMIC}$/$Dap160^{RNAi}$ | <br>9 NMJs/4 larvae<br>10 NMJs/5 larvae | <br>One step association curve<br>Mann-Whitney U test on taus |

| **Figure/ Experiment** | **Genotype/Conditions** | **N** | **Statistical Test(s)** |
|---|---|---|---|
| 5 S1A-B<br>Nwk$^{1-633}$<br>WASp<br>Dap160$^{CD}$ | DOPC/DOPE/DOPS/PI(4,5)P$_2$ = 75/15/5/5<br>1μM<br>2 μM<br>3μM | 3 tech replicates per group | ANOVA+<br>Tukey's multiple comparison test |
| 5 S1C<br>Nwk$^{1-731}$<br>WASp<br>Dap160$^{CD}$ | DOPC/DOPE/DOPS/PI(4,5)P$_2$ = 70/15/5/10<br>1μM<br>1μM<br>3μM | 3 tech replicates per group | ANOVA+<br>Tukey's multiple comparison test |
| 5 S1D<br>Nwk$^{1-754}$<br>WASp<br>Dap160$^{SH3CD}$ | DOPC/DOPE/DOPS/PI(4,5)P$_2$ = 70/15/5/10<br>250nM<br>1μM<br>250nM5% PI(4,5)P$_2$ GUVs | Representative from:<br>5 GUVs imaged<br>10 GUVs imaged | NA |
| 6A-C<br>Control<br>*nwk*$^{1/2}$ | <br>C155-GAL4/+ or Y; UAS-GMA/+<br>C155-GAL4/+ or Y; UAS-GMA, *nwk*$^{2}$,*h*/ *nwk*$^{1}$ | <br>1606 patches/20 NMJ/8 larvae<br>1928 patches/22 NMJ/8 larvae | <br>Mann-Whitney (B)<br>Kolm.-Smirnoff (C) |
| 6D-F<br>FL<br>ΔCD | C155-GAL4/+; dap160$^{\Delta1}$/Df3450; UAS-GMA/X<br>X=UAS-Dap$^{FL}$::mCherry/+<br>X=UAS-Dap$^{\Delta CD}$/+ | <br>1279 patches/14 NMJ/7 larvae<br>1937 patches/17 NMJ/7 larvae | <br>Mann-Whitney (E)<br>Kolm.-Smirnoff (F) |
| 6 S1A-C | Same data as 6A-C | As 6A-C | t-test (B)<br>t-test (C) |
| 7B-C | C155-GAL4/+; UAS-lifeact::Ruby/AP2::GFP | 14 NMJ/3 larvae | NA |
| 7D-E<br>Control<br>*shi*TS1 | <br>C155-GAL4/Y; UAS-GMA/+<br>C155-GAL4, shiTS1/Y; UAS-GMA/+ | <br>669 patches/16 NMJ/8 larvae<br>901 patches/19 NMJ/8 larvae | <br>Welch's t test (D) |
| 7A-BF-G<br>Control<br>*nwk*$^{1/2}$<br>*shi*$^{TS1}$ | <br>C155-GAL4/Y; UAS-GFP/+<br>C155-GAL4/Y; UAS-GFP/+; *nwk*$^{1}$/*nwk*$^{2}$*h*<br>C155-GAL4, shi$^{TS1}$/Y; UAS-GFP/+ | <br>23 NMJ/4 larvae<br>24 NMJ/4 larvae<br>23 NMJ/4 larvae | Kruskal-Wallis +<br>Dunn's multiple comparison test |
| 7C-DH-I<br>Dap$^{FL}$<br>Dap$^{\Delta D}$<br>Dap$^{\Delta CD}$<br>*shi*$^{TS!}$ | rescues= C155-GAL4/+; dap160$^{\Delta1}$/Df3450; X<br>X=UAS-Dap$^{FL}$::mCherry/+<br>X=UAS-Dap$^{\Delta D}$/+<br>X=UAS-Dap$^{\Delta CD}$/+<br>C155, shiTS1/Y x UAS-RFP | <br>32 NMJs/8 larvae<br>12 NMJs/3 larvae<br>12 NMJs/3 larvae<br>8 NMJs/2 larvae | Unpaired t-tests to dish-matched controls |
| 7 S1A-C<br>Control<br>*nwk*$^{1/2}$ | <br>C155-GAL4/Y; UAS-GFP/+<br>C155-GAL4/Y; UAS-GFP/+; *nwk*$^{1}$/*nwk*$^{2}$*h* | <br>15 NMJ/4 larvae<br>19 NMJ/5 larvae | <br>Mann-Whitney (B)<br>t-test (C) |
| 7 S1D-F<br>Dap$^{FL}$<br>Dap$^{\Delta D}$<br>Dap$^{\Delta CD}$ | rescues= C155-GAL4/+; dap160$^{\Delta1}$/Df3450; X<br>X=UAS-Dap$^{FL}$::mCherry/+<br>X=UAS-Dap$^{\Delta D}$/+<br>X=UAS-Dap$^{\Delta CD}$/+ | <br>16 NMJs/4 larvae<br>15 NMJs/4 larvae<br>16 NMJs/4 larvae | <br>ANOVA+Tukey's multiple comparison test |
| Figure 7S2A-B<br>DapRNAi<br>mCh<br>Dap$^{FL}$<br>Dap$^{\Delta D}$<br>Dap$^{\Delta CD}$ | <br>C155-GAL4, UAS-Dcr2/Y; UAS Dap160-RNAi/+<br>C155-GAL4/+; UAS-CD8::RFP/+<br>X=UAS-Dap$^{FL}$::mCherry/+<br>X=UAS-Dap$^{\Delta D}$/+<br>X=UAS-Dap$^{\Delta CD}$/+ | <br>24 NMJ/6 larvae<br>22 NMJ/6 larvae<br>21 NMJ/6 larvae<br>16 NMJ/5 larvae<br>23 NMJ/6 larvae | <br>ANOVA+Tukey's multiple comparison test |
